# Evolutionary dynamics of the *tgr* gene family in *Dictyostelium* allows escape from Crozier’s Paradox

**DOI:** 10.1101/2025.10.22.684067

**Authors:** Rafael A. Polo Prieto, Sean McFadyen, Fabiola Medina, Mische Holland, Cathleen M.E. Broersma, Jenna Masterson, Courtney Masterson, Hui Hui Phua, Jennie Kuzdzal-Fick, Murray P. Cox, Tera C. Levin, Elizabeth A. Ostrowski

**Author notes:** Contributed equally (Rafael A. Polo Prieto, Sean McFadyen).

## Abstract

Crozier’s Paradox states that genetic kin recognition will select for its own demise, and theoretical analyses over the past few decades have supported this conclusion. However, empirical evidence is far more limited. Here we examine support for Crozier’s Paradox in two kin recognition genes in the social amoeba *Dictyostelium discoideum*, *tgrB1* and *tgrC1*, which enable cells to recognize and reject unrelated cells during cooperative multicellular development. We find that co-isolated amoebae, taken from just a few grams of soil, harbor highly diverged recognition alleles, indicating plentiful sequence variation for social decision-making on a micro-scale. *tgrB1* and *tgrC1* belong to a large gene family, whose members vary in their levels of polymorphism, strength of selection, and retention of polymorphism. We do not see strong evidence for the paradox, in the form of positive frequency-dependent selection, or its outcome, in the form of depleted genetic variation. Consistent with previous findings, we detect recombination among the duplicated *tgrB* and *tgrC* genes, which suggests a mechanism to generate novel alleles and replenish genetic variation. We suggest that the evolutionary dynamics of large gene families may be essential for rapidly generating novel variation that can prevent Crozier’s Paradox from taking effect.

## Introduction

Cooperative groups often involve costs that are shared unequally among the members. Such systems are prone to the emergence of selfish individuals, who access the benefits of membership without paying its full costs (Sachs et al. 2004; West et al. 2006; Ghoul et al. 2014). One way in which selfishness can be defeated is if cooperative benefits are preferentially directed to kin, ensuring that one’s own alleles (especially those for the cooperative act itself) benefit from any sacrifices made (Hamilton 1964a; Hamilton 1964b; Buss 1982; Gadagkar 1985; Agrawal 2001; Boomsma and d’Ettorre 2013; Sheehan et al. 2017). The ability to distinguish kin from non-kin is thus an essential component of stable, complex cooperation.

Despite the potential benefits of genetic kin recognition, evolutionary theory indicates that these systems are prone to depleting the very genetic variation on which they rely (Crozier 1986; Grafen 1990; Blaustein et al. 1991; Gardner and West 2007; Rousset and Roze 2007; Field et al. 2018). The problem was first described nearly a half century ago by Crozier, who pointed out that rare alleles provide the most accurate information about relatedness, but individuals with common alleles can engage in more cooperation and thus receive a disproportionate share of its benefits (Crozier 1986). This dilemma, known as Crozier’s Paradox, predicts that common alleles should fix through positive frequency-dependent selection, leading to loss of genetic diversity and therefore loss of the functional system (Crozier 1986; Gardner and West 2007; Rousset and Roze 2007; Holman et al. 2013; Field et al. 2018).

Crozier’s original paper consisted of a simple model: the entire paper, including references, was just two pages long. However, Rousset and Roze (2007) conducted a more thorough analysis of the problem, concluding that the Crozier effect occurs over a wide range of parameter values. They also reviewed what was known at the time about the genetic bases of kin recognition. They concluded that Crozier’s intuition that kin recognition must be maintained by selection for ‘something else’—with kin recognition as a secondary benefit— could well be right (Rousset and Roze 2007).

Renewed interest in Crozier’s Paradox has led to additional ideas about how it could be prevented (reviewed in Green et al. 2024). Given the dual roles of the major histocompatibility (MHC) loci in both kin recognition and parasite avoidance, the ‘something else’ is typically believed to be pathogen recognition (Scott et al. 2023). However, the limited number of well-characterized, non-MHC kin recognition systems makes it difficult to know if the MHC’s dual roles reflect happenstance or necessity. Bastiaans et al. (2015) showed that Crozier’s Paradox arises in fungi, which have vegetative incompatibility (VIC) loci that govern hyphal fusion/rejection, unrelated to the MHC of vertebrates (Glass et al. 2000; Aanen et al. 2010; Arshed et al. 2023). They proposed that the benefit of having common alleles could be counter-balanced by an increased risk of acquiring somatic parasites, although they did not test the idea. Holman et al. (2013) proposed that the primary role of recognition genes could be disassortative mating, noting that some kin recognition genes, including VICs, also function in mate recognition. Finally, Scott et al. (2022) proposed a general solution to Crozier’s Paradox that does not rely on secondary functions at all. They showed that if individuals can ‘prospect’ before cooperating (Green et al. 2024), then those with rare alleles can find one another. So long as the search costs for these rare types are low, the process can prevent Crozier’s Paradox from being triggered.

The social amoeba *Dictyostelium discoideum* is one of the best-known examples of a species with genetic kin recognition (Hirose et al. 2011; Ostrowski 2019). During its unicellular stage, these haploid amoebae prey upon microbes and divide asexually to produce daughter cells. Upon starvation, however, aggregate and develop into a motile, multicellular slug and later a multicellular fruiting body. Some cells, having formed the upright stalk of the fruiting body, vacuolize and die. Others form resistant spores, which survive atop the stalk and disperse. Multicellular development is thus cooperative, and stalk formation is altruistic: stalk cells die to benefit the survival and dispersal of the spore cells (Strassmann et al. 2000; Ostrowski 2019).

*Dictyostelium*’s aggregative multicellularity poses a conundrum. Because genetically different strains can co-aggregate, selection can favor cells that behave selfishly, preferentially forming spores and not stalk (Strassmann et al. 2000). However, *Dictyostelium* can also recognize and reject foreign cells from its aggregations (Ostrowski et al. 2008). Cells of different genotypes initially co-aggregate but separate into different structures as development proceeds. Although the separation is imperfect, at least in laboratory settings, it increases the relatedness of strains that share the spoils of multicellular development—that is, the stalk (Ostrowski et al. 2008). Thus, genetic kin recognition potentially stabilizes altruistic cooperation in this species (Ostrowski 2019).

In *Dictyostelium*, kin recognition is carried out by two cell-surface proteins, TgrB1 (the ligand) and TgrC1 (the receptor), which bind one another following aggregation (Hirose et al. 2011). When binding occurs, the cells remain together during multicellular development, resulting in chimeric multicellularity. The genes are named for their protein domains: **<u>T</u>**ransmembrane, IPT/**I**mmuno**<u>g</u>**lobulin-like **<u>E</u>**-set **<u>R</u>**epeat domains, or *tiger* (*tgr)* genes. The functions of the TgrB1 and TgrC1, in both allorecognition and multicellular development, have been well-studied (Benabentos et al. 2009; Hirose et al. 2011; Hirose et al. 2015; Hirose et al. 2017; Katoh-Kurasawa et al. 2024). They not only govern rejection of foreign cells, but also influence the adoption of the spore versus stalk cell fate, thus directly affecting altruism in response to relatedness (Kuzdzal-Fick et al. 2023; Katoh-Kurasawa et al. 2024).

Crucially, several features of *Dictyostelium* biology make it unlikely that the aforementioned solutions to Crozier’s Paradox apply to this case. First, its two kin recognition genes do not have any other established recognition functions. So far, *tgrB1* and *tgrC1* are only known to be expressed in a short two-hour developmental window when cells of different genotypes sort out and adopt their initial cell fates (Benabentos et al. 2009; Hirose et al. 2011; Ho and Shaulsky 2015). This restricted period of expression makes it unlikely that genetic variation can be maintained by selection on secondary functions, such as pathogen recognition. The genes also do not govern mate recognition. The mating type locus has been identified (Bloomfield et al. 2010; Bloomfield 2011); it is not even on the same chromosome, so even linkage effects can be ruled out. ‘Prospecting before cooperating’ (Scott et al. 2022) is also not an option: *Dictyostelium* uses a generic chemoattractant, cyclic AMP, that allows even other dictyostelid species to join the aggregates (Raper and Thom 1941). While they do sort out after aggregation, search costs are probably high: even if rare allotypes locate one another in a sea of other cells, they will likely lack the quorum needed to form their own multicellular structure. An inability to identify ‘like’ individuals prior to joining the group, combined with high costs of being rare, means that rare allotypes are likely disadvantaged relative to their common counterparts. For these reasons, whether *Dictyostelium* experiences Crozier’s Paradox—and if so, how it evades its effects—remains a mystery.

Here we examine the level and retention of polymorphism in *Dictyostelium tgr* genes and assess the applicability of Crozier’s Paradox to its kin recognition system. To do so, we sampled co-occurring natural isolates from small soil samples. This approach revealed high diversity in kin recognition genes at the local scales most relevant to social interactions. We also assessed the molecular evolution of these genes across multiple sites to test the generality of our conclusions about the level of variation and form of selection acting on these genes in nature. Finally, we used long-read Oxford Nanopore technology to sequence and assemble whole genomes, which allowed us to characterize the gene family comprehensively. Our results reveal a structurally complex locus, reminiscent of the MHC locus in vertebrates, which can generate new alleles in part through intra- and inter-locus recombination. Despite high inputs of genetic variation, likely effective in preventing Crozier’s Paradox, we nevertheless detect strong positive or diversifying selection on only a subset of the *tgrBC* genes, supporting a dominant role for *tgrB1* and *tgrC1*.

## Results

### The *tgr* Gene Family

The *tgrB1* and *tgrC1* kin recognition genes belong to a large gene family, which consists of an estimated 63 members in the AX4 reference genome (Fig. 1A), making it one of the largest gene families in *Dictyostelium*. Like *tgrB1* and *tgrC1*, other *tgr* genes primarily show the same sequence of domains, although some lack the transmembrane domain, such as *tgrB2* (Fig. 1B). The canonical genes, *tgrB1* and *tgrC1*, are found in a head-to-head arrangement (Fig. 1C; Benabentos et al. 2009). Other *tgr* genes are also paired (examples in Fig. 1C). Gene duplications have resulted in clusters of *tgr* genes with intervening non-*tgr* genes <u>(red bars;</u> Fig. 1A). While some clusters consist primarily of duplicated head-to-head pairs, others consist of unpaired genes (e.g., *tgrA*) or a mix of paired and unpaired genes. Others occur as isolated pairs that lack nearby duplicates (e.g., *tgrD1* and *tgrE1*). Others still are singletons with respect to their distance to other *tgr* genes on the chromosome (e.g., *tgrR1*).

**Figure 1.**
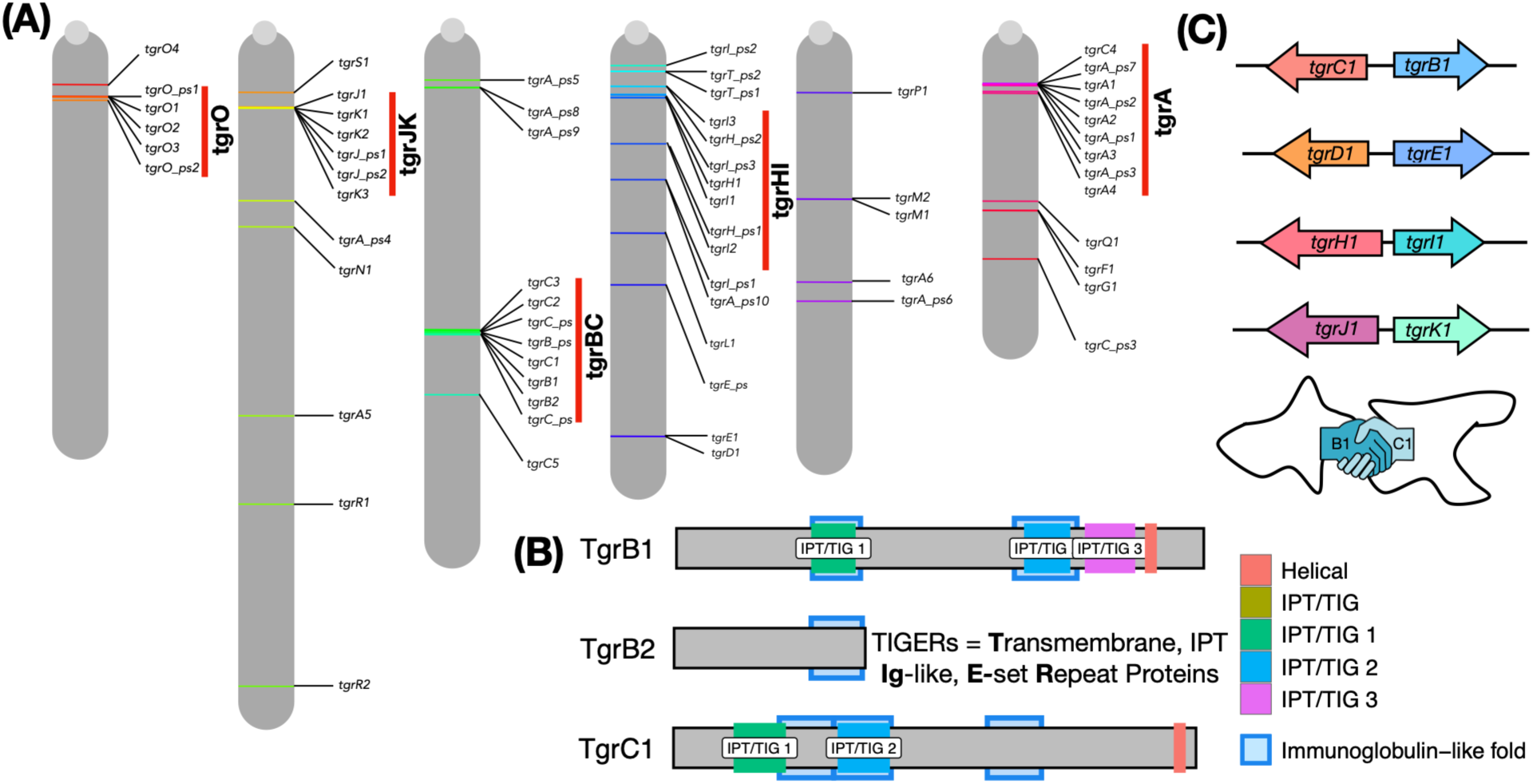
The *tgr* gene family in *Dictyostelium discoideum*. **(A)** Chromosomal locations of *tgr* genes annotated in the *D. discoideum* lab strain AX4, according to the GFF available from dictybase.org. Clusters, indicated by vertical red bars, consist of duplicated genes and pseudogenes in proximity on the chromosome. **(B)** Protein domains present in TgrB1, TgrB2, and TgrC1; additional proteins are shown in Fig. S1. Filled boxes indicate Uniprot IPT/TIG or Ig E-set repeat domains (Interpro IDs IPR002909, IPR014756), whereas shaded blue boxes indicate the homologous superfamily Ig-like folds (IPR013783). **(C)** Examples of *tgr* genes that sit in head-to-head pairs. In *tgrB1* and *tgrC1*, the region between genes pairs harbors a shared promoter, which enables their coordinated transcription at around 9 hours of development, when initial cell-type differentiation takes place and co-aggregated genotypes separate.

As a whole, the *tgr* gene family is an exemplar of ‘birth-and-death’ evolution (Nei et al. 1997), a common feature of recognition genes (Nei et al. 1997; Piontkivska and Nei 2003; Karev et al. 2006). Specifically, they form clusters of closely related genes (frequent ‘birth’ through gene duplication), with many apparent pseudogenes (frequent ‘death’ through inactivating mutations). Owing to their rapid evolution, however, it is unknown whether loci that are pseudogenes in the reference strain encode functional genes in other strains, or vice versa. Existing transcriptome data also indicate low or undetectable expression for most members of the gene family (Fig. S2), at least under conditions tested so far, making it challenging to distinguish functional from nonfunctional copies.

### Illumina sequencing is inaccurate for the *tgr* gene family

We initially compared pre-existing Illumina sequences (Ostrowski et al. 2015) to Sanger sequences (Benabentos et al. 2009) for the same strains and found many sequence differences (strain list and sequencing methodologies provided in Table S1). We also sequenced some strains with the newer Illumina technology, which used 150-bp paired end reads (see Supplement for details of sequencing, mapping, and SNP calls). Simultaneously, we used gold-standard Sanger sequencing to sequence six loci: *tgrB1*, *tgrC1*, as well as *tgrB2*, *tgrC2*, *tgrC3*, and *tgrC5*. These results were largely consistent with prior Sanger sequences for these strains (Benabentos et al. 2009) but again did not match the Illumina sequences, despite updated filtering and SNP calling methods, suggesting that the sequences derived from short-reads were inaccurate (Fig. 2).

**Figure 2.**
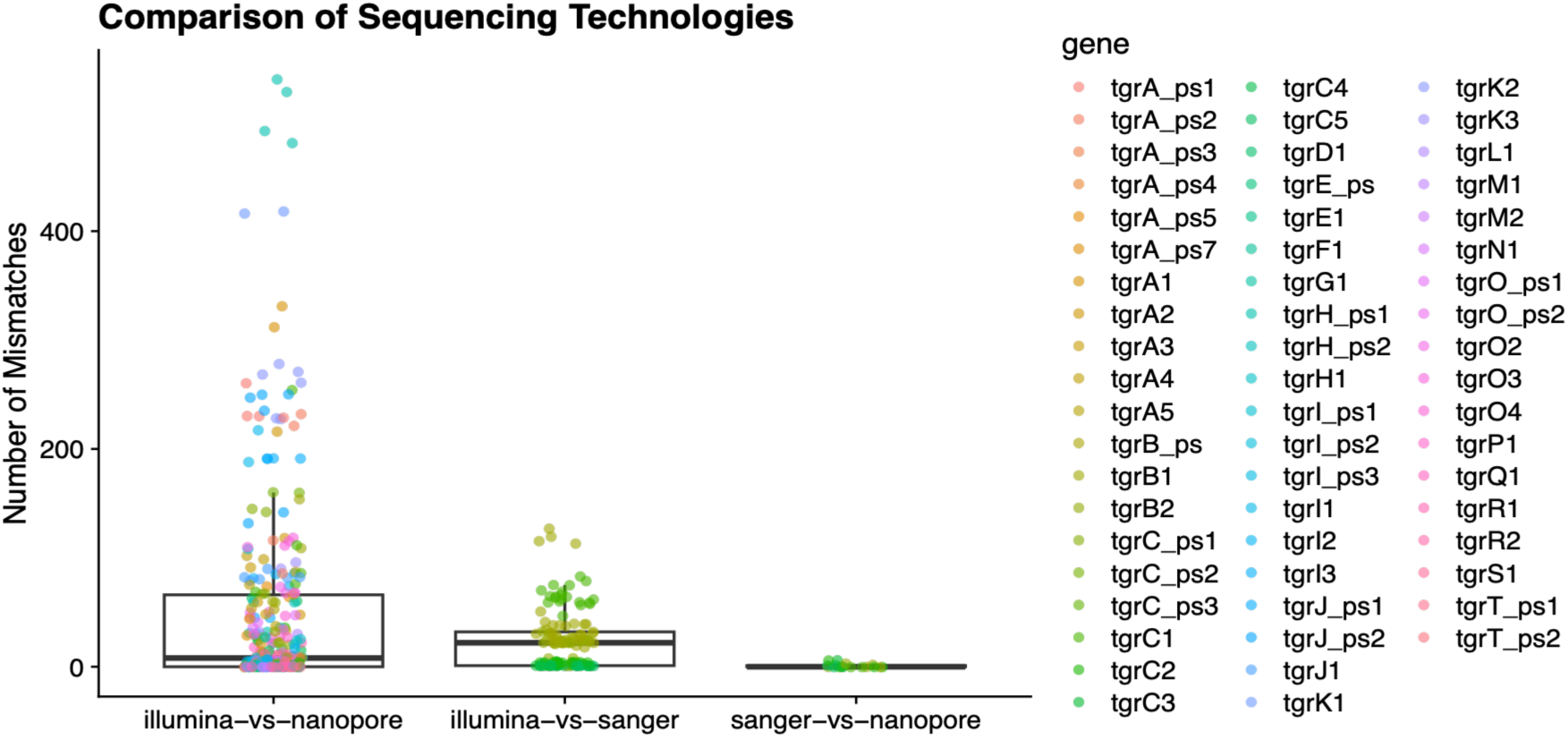
Nucleotide mismatches between Sanger, Illumina, and Nanopore sequencing technologies. Each point represents a comparison of the same strain’s sequence, for the same gene, as determined by the two methods listed on the x-axis. Illumina and Nanopore methods provided sequences for all *tgr* genes (colored dots) whereas Sanger sequencing was only performed for *tgrB* and *tgrC* genes. Nanopore and Sanger sequences are nearly identical to each other, but both differ from Illumina sequences, suggesting that short-read based approaches do not work well for this large polymorphic gene family.

### Nanopore and Sanger sequencing for a given strain produce near identical sequences, suggesting both technologies are highly accurate

Owing to the numerous closely related gene paralogs in the *tgr* gene family, high polymorphism in these genes, and the low complexity of the *D. discoideum* genome (Eichinger et al. 2005), we surmised that these complex regions would benefit from long-read sequencing. We also suspected the existence of structural variation. For example, we were unable to PCR-amplify *tgrB2* in nearly half of the strains from Texas, suggesting polymorphic structural variation within populations. Amplification using primers placed in surrounding genes eventually revealed a 2-kb deletion in the region where *tgrB2* is expected (Fig. S3). For all of these reasons, we undertook Nanopore genome sequencing and *de novo* assembly of 9 strains (Table 1; strain details in Table S1). Three of these strains potentially lacked *tgrB2*, so long-read sequencing would confirm this putative structural variation. Two other strains were suspected cryptic species based on earlier Illumina sequencing (see “Outgroup strains” in Methods). High-quality assemblies for a closely related outgroup would be beneficial for molecular evolution analyses, as divergence to the *D. citrinum* outgroup used previously was too great (Ostrowski et al. 2015).

**Table 1.** Assembly statistics for nine *Dictyostelium* strains sequenced using Nanopore long-read sequencing.

| Species Status | Strain | #Contigs | Genome Size | GC% | N50 | L50 | BUSCO | Coverage |
| --- | --- | --- | --- | --- | --- | --- | --- | --- |
| *Dd* | EO617 | 66 | 35,579,755 | 23.38 | 4,852,304 | 3 | 92.9 | 103 |
| Cryptic | EO1089 | 213 | 39,873,666 | 27.35 | 1,062,780 | 11 | 94.1 | 130 |
| Cryptic | EO1176 | 71 | 39,745,864 | 27.32 | 3,515,978 | 5 | 93.4 | 44 |
| *Dd* | EO1224 | 203 | 33,836,964 | 22.55 | 2,858,045 | 5 | 93.8 | 94 |
| *Dd* | EO1236 | 84 | 33,748,730 | 22.63 | 4,102,943 | 4 | 93 | 47 |
| *Dd* | EO1271 | 96 | 33,819,639 | 22.7 | 3,014,741 | 5 | 90.6 | 21 |
| *Dd* | QS18 | 116 | 33,911,281 | 22.61 | 2,417,048 | 6 | 92.5 | 150 |
| *Dd* | QS33 | 87 | 33,866,725 | 22.62 | 3,021,021 | 4 | 93.3 | 245 |
| *Dd* | QS68 | 107 | 35,345,620 | 23.35 | 4,348,941 | 4 | 92.9 | 110 |

**Table 2.** McDonald-Kreitman test components, neutrality index (NI), alpha (percentage of adaptive substitutions), and direction-of-selection (DoS) results for three genes. The test was significant for three tgr genes, *tgrB1*, *tgrC1*, and *tgrC4*. For comparison, we include the results for *tgrB2*, despite not being significant.

| gene | Dn | Ds | Pn | Ps | NI | alpha | DoS | adjusted *P* |
| --- | --- | --- | --- | --- | --- | --- | --- | --- |
| *tgrC1* | 164 | 41 | 215 | 123 | 0.44 | 0.56 | 0.16 | 0.001** |
| *tgrC4* | 55 | 81 | 11 | 2 | 8.10 | -7.10 | -0.44 | 0.04* |
| *tgrB1* | 22 | 26 | 179 | 84 | 2.52 | -1.52 | -0.22 | 0.04* |
| *tgrB2* | 53 | 21 | 38 | 10 | 1.51 | -0.51 | -0.08 | 0.74 (NS) |

Assembly of Nanopore long-reads resulted in high-quality assemblies, with >90% BUSCO coverage (Table 1). After annotation, we incorporated our sequences into the existing *D. discoideum* phylogeny (Holland, Ahmed, Young, Drurey, et al. 2025), which confirmed the placement of our two suspected cryptic species strains, EO1089 and EO1176, in a sister clade to the main *D. discoideum* clade (Fig. 3). The Nanopore sequences also closely matched those generated by Sanger sequencing (Fig. 2). Longer reads, combined with *de novo* assembly, likely circumvented the mis-mapping of short reads caused by numerous closely related gene paralogs and high levels of polymorphism.

**Figure 3.**
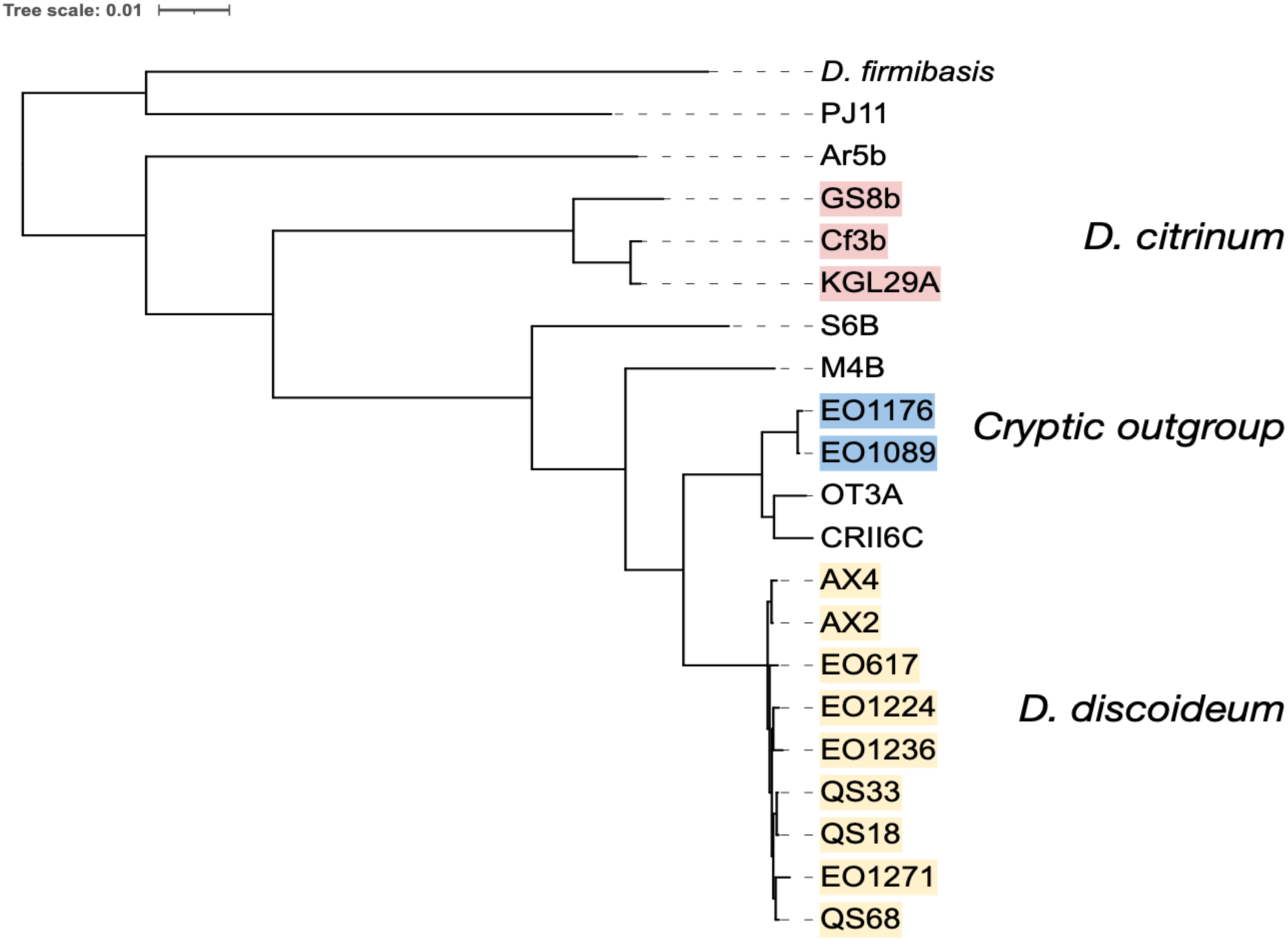
Protein-based phylogenetic tree of assembled strains. Strains were incorporated into the published phylogeny of Holland et al. (2025) of 174 conserved BUSCO genes to confirm species designations. We designate EO1089 and EO1176 (in blue) as a cryptic outgroup, because they co-occurred in soil samples with *D. discoideum* strains, were highly genetically diverged based on both Illumina and Nanopore sequencing, but they could not be distinguished morphologically from *D. discoideum*. See Methods for further discussion.

### Identifying *tgr* genes in genome assemblies

We first analyzed the presence and absence of the gene family members using Liftoff, which lifts over gene annotations among assemblies from the same or closely related species by aligning genes (specifically, exons) rather than entire genomes (Shumate and Salzberg 2021). These analyses successfully identified many *tgr* gene family members, and putative absences were typically observed across multiple strains (Fig. 4). Manual inspection of genes designated as *tgr* pseudogenes in AX4 revealed a variety of patterns: many of these alleles appear to be pseudogenes, but there were also exceptions where one or a few strains had uninterrupted reading frames over most or all of the gene.

**Figure 4.**
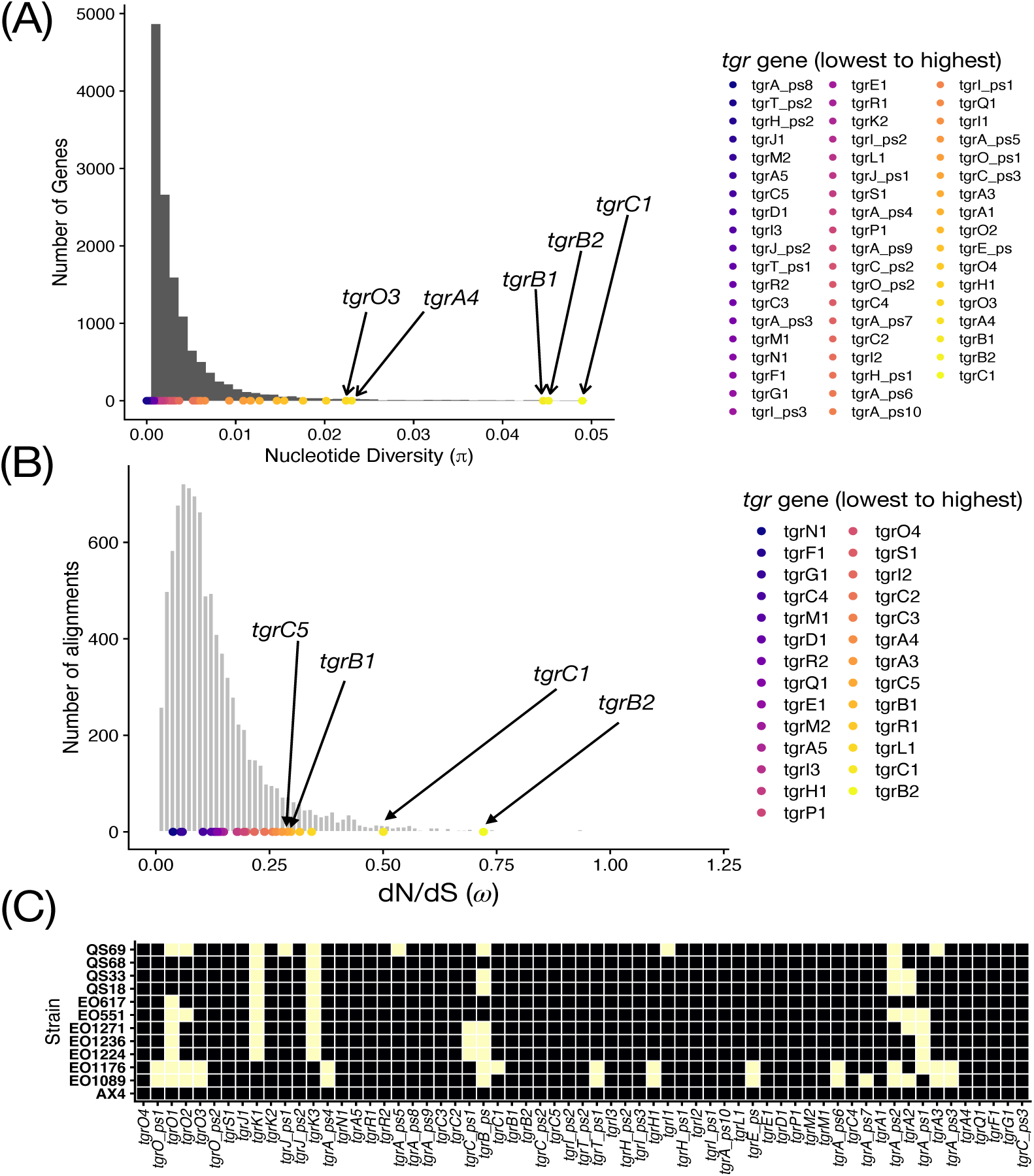
**(A)** Nanopore genome assemblies reveal the *tgr* gene family is highly polymorphic with *tgrB1*, *tgrB2*, and *tgrC1* showing the greatest levels of nucleotide variation. Gray bars indicate the distribution of nucleotide diversity for all genes in the genome, and circles indicate members of the *tgr* gene family. **(B)** Genome-wide ratio of the rates of nonsynonymous (dN) to synonymous (dS) substitution (*_ω_*), an indicator of positive or diversifying selection. Each value represents the mean of all pairwise comparisons between a *D. discoideum* strain and one of the cryptic outgroup strains. The *tgrB2* gene is ranked in the 99th percentile of the genome-wide distribution, whereas *tgrC5* and *tgrB1* both reside in the 92nd percentile. **(C)** Variation in *tgr* gene family content among strains. The x-axis shows each *tgr* gene, based on the reference genome annotation, with ‘ps’ indicating a pseudogene in the reference genome. Genes are ordered according to chromosome (one to six) and then by position. The Y-axis indicates each strain. Black boxes indicate gene presence based on Liftoff analysis, whereas yellow boxes indicate gene absence.

To further assess the composition of the gene family, we used tblastn with the AX4 *tgr* proteins as queries to search the assemblies for partial or complete *tgr* sequences. Overall, we identified more than 75 genes or fragments per strain that were significant hits to the *tgr* gene family (Fig. 5). We also observed ∼3x more *tgrC*-like (ligand) sequences than *tgrB*-like (receptor) sequences, an asymmetry that is also present in the AX4 reference genome (8 *tgrC* vs 3 *tgrB*). Filtering our *tgrB*/*C* sequences to those where the *tgrB* and *tgrC* genes are in a head-to-head arrangement (indicative of a functional receptor-ligand pair), we found only one to two *BC* pairs per strain, which correspond well to the positions of *tgrB1*-*tgrC1* and *tgrB_ps* and *tgrC_ps1* in the AX4 reference genome. A third fragmentary pair is present, but not in a head-to-head arrangement. Regardless, there are many *tgr* gene sequences per strain (>75 genes or gene fragments) and many paired head-to-head pairs of *tgr* genes (>20) per genome, indicating a large pool of *tgr*-like sequences.

**Figure 5.**
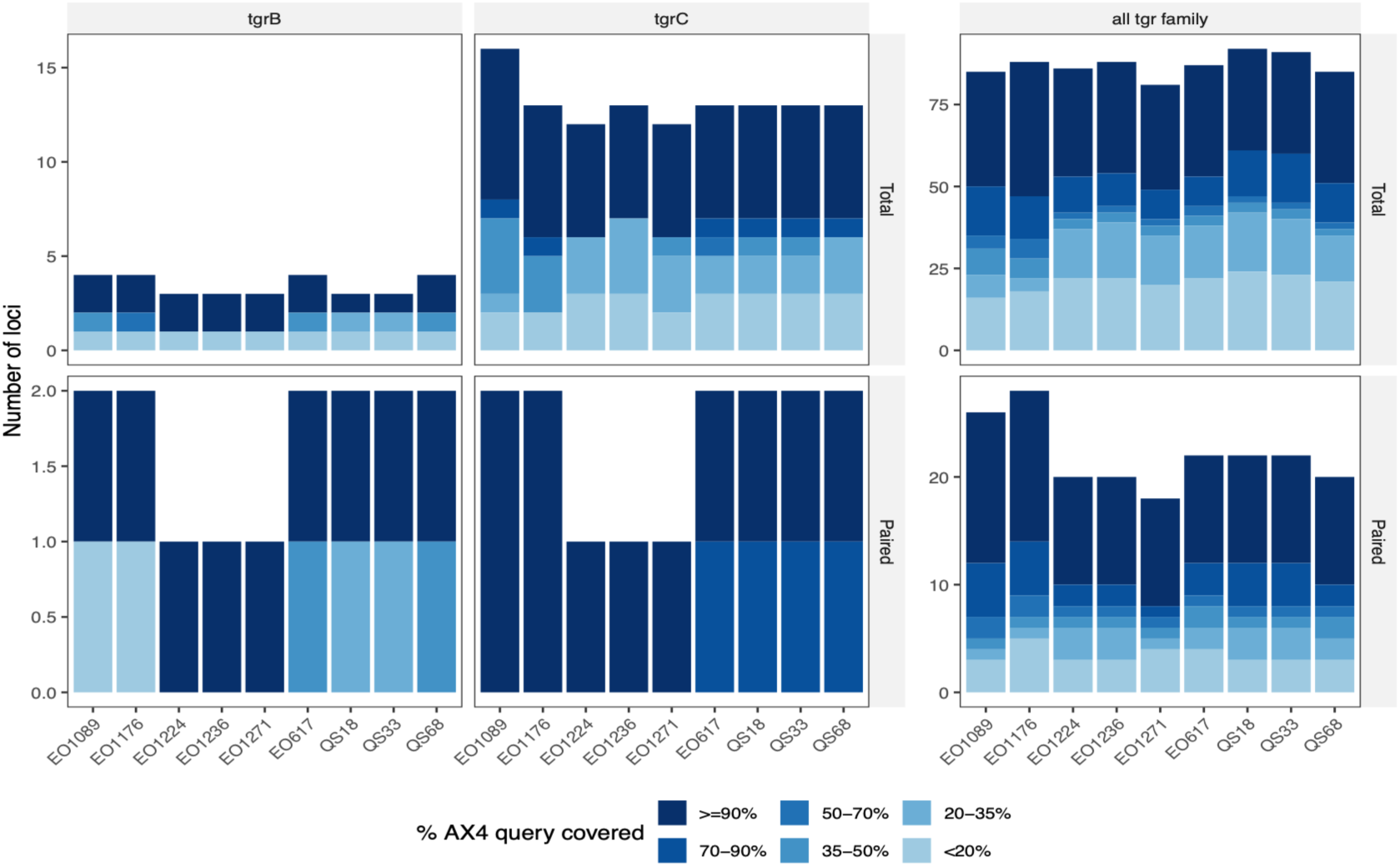
Full-length and fragmentary *tgr* genes in the genome assemblies. We used tblastn and AX4 protein queries to find significant hits to *tgr* genes. (A) Top row: number of *tgrB*, *tgrC,* or total *tgr* sequences, colored according to the percentage of the query (i.e., AX4 sequence) they cover. Bottom row: number of *tgrB*, *tgrC,* or total *tgr* blast hits occurring in head-to-head pairs.

Despite high-quality assemblies, regions surrounding the *tgrBC* locus did not align well (Fig. S4). Contigs harboring the *tgrBC* locus, while aligning well over much of their length (at least between pairs of the bona fide *D. discoideum* strains), typically contained multiple unalignable regions around the *tgrBC* locus, even with relaxed mapping parameters allowing up to 20% sequence divergence (Fig. S5). A few closely related strain pairs did produce perfect (i.e., contiguous) alignments through the *tgrBC* locus from independent assemblies (for example, Fig. S4B), so we are confident that this pattern is caused by sequence divergence and structural variation rather than misassembly.

### *tgrB1* and *tgrC1* — but also *tgrB2* — are extraordinarily polymorphic and rapidly evolving

To assess *tgr* gene variation compared to the genomic background, we aligned the sequences of each gene in the genome using our Liftoff-generated gene annotations. We observed extremely low sequence variation on average in this species, with a mean and median nucleotide diversity across all *D. discoideum* genes of 0.003 and 0.0014, slightly higher than previous estimates of 0.001 (Ostrowski et al. 2015). It was therefore striking that the regions surrounding the *tgrBC* locus exceed 20% nucleotide divergence—precluding alignment altogether—in strain pairs that are, on average, 99.7% identity genomewide. These results indicate the extreme departure of the *tgrBC* locus from the rest of the genomic background.

Having identified most *tgr* genes, we could differentiate the orthologs from the paralogs and investigate selection on individual *tgr* genes. For nucleotide diversity, we observe *tgr* genes in the right tail of the distribution, with genes from the *tgrBC* cluster dominating these outliers (Fig. 4A). In addition, the designated *tgr* pseudogenes do not dominate the tail of the distribution, suggesting that the unusual level of variation is not an artefact of the many pseudogenes. Instead, the functional genes *tgrB1*, *tgrB2*, and *tgrC1* are amongst the most polymorphic genes in the genome, all exceeding the 99.6^th^ percentile for polymorphism and nearly one-quarter (24%) of *tgr* genes exceeding the 95^th^ percentile.

To assess evidence of selection on *tgrBC* genes, we examined the ratio of the rates of nonsynonymous to synonymous substitution (dN/dS), using the average dN/dS of all pairs of *D. discoideum* strains and their cryptic outgroup strains (EO1089 and EO1176). These analyses place *tgrB1, tgrC1, tgrB2,* and some other *tgr* genes in the tail of the distribution (99^th^, 97^th^, and 92^nd^ percentile of the genome-wide distribution; Fig. 4B). Sliding window analysis of dN/dS furthermore indicates that positive selection is limited to certain regions of the genes (Fig. 6). In *tgrB1* especially, dN/dS is extreme (>7) at the start of the gene, whereas the remainder of the gene shows dN/dS<1, consistent with purifying selection.

**Figure 6.**
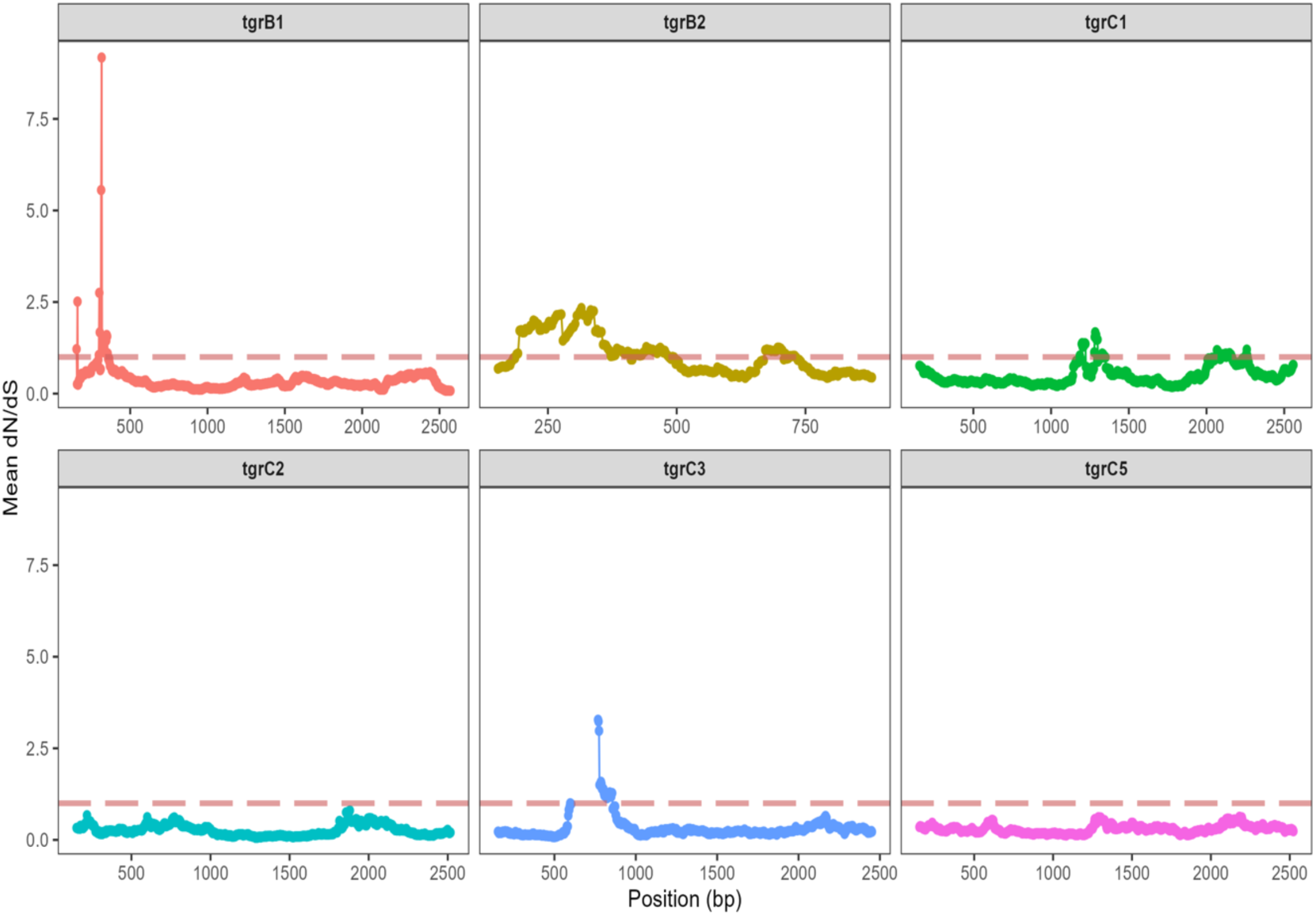
Mean dN/dS across all pairwise comparisons between *D. discoideum* and each of the two cryptic outgroup strains, EO1089 and EO1176. Dotted line indicates a dN/dS ratio of one. Elevations above one mean that nonsynonymous substitutions accumulate at a faster rate than synonymous substitutions, an indicator of positive selection.

To verify mixed modes of selection acting on *tgrB1*, we used PAML to test for positive selection, as well as variable strength of selection across the gene. Likelihood ratio tests, comparing models with or without positive or site-specific selection, indicate that *tgrB1*, *tgrC1*, and *tgrB2* all experience positive selection that is variable in strength across sites (Table S2). However, the models also estimate that only 0.6% of *tgrB2* sites are experiencing positive selection – which works out to only 6 or 7 sites, given the size of the gene (Table S3).

Finally, we used McDonald-Kreitman (MK) tests to compare the ratio of within-species nonsynonymous to synonymous polymorphisms (Pn/Ps) to the ratio of between-species nonsynonymous to synonymous fixed differences (Dn/Ds). Under neutrality, Pn/Ps is expected to equal Dn/Ds. Among the *tgr* genes, three—*tgrB1*, *tgrC1*, and *tgrC4*—had significant Fisher’s Exact tests after correction for multiple testing across the *tgr* gene set. Their p-values also fell within the lowest 5% of the genome-wide distribution, based on 9,637 genes for which the test could be computed. These results indicate patterns of nonsynonymous polymorphism and divergence that are unusual relative to other genes in the genome.

Surprisingly, *tgrC1* and *tgrB1* departed from neutrality in opposite directions. *tgrC1* showed an excess of nonsynonymous fixed differences relative to polymorphisms (Dn/Ds > Pn/Ps), whereas *tgrB1* showed an excess of nonsynonymous polymorphisms relative to fixed differences (Pn/Ps > Dn/Ds). For *tgrB1*, this pattern is reflected in a neutrality index greater than one (NI=2.52), which can result from balancing selection or purifying selection against nonsynonymous variants (Fijarczyk and Babik 2015). Given the spatial variation in dN/dS across *tgrB1*, together with its negative Tajima’s D (described below), we interpret these results as reflecting heterogeneous selection across the gene, with positive or balancing selection in some regions and purifying selection in others. *tgrC4* also showed an excess of nonsynonymous polymorphism relative to divergence (NI=8.10), but its functional significance is less clear because its expression and biological role have not been established.

### Spatial analyses indicate plentiful genetic variation at the centimeter-scale in nature

Escaping Crozier’s Paradox requires more than just polymorphism in the species as a whole: variation must be present locally, over spatial scales where individuals assess potential social partners and decide whether to cooperate with them. We thus focused on detecting selection at the centimeter-scale in four natural populations (Figure 7). The sites varied in pairwise distance from ∼100 m to >1,000 km.

**Figure 7.**
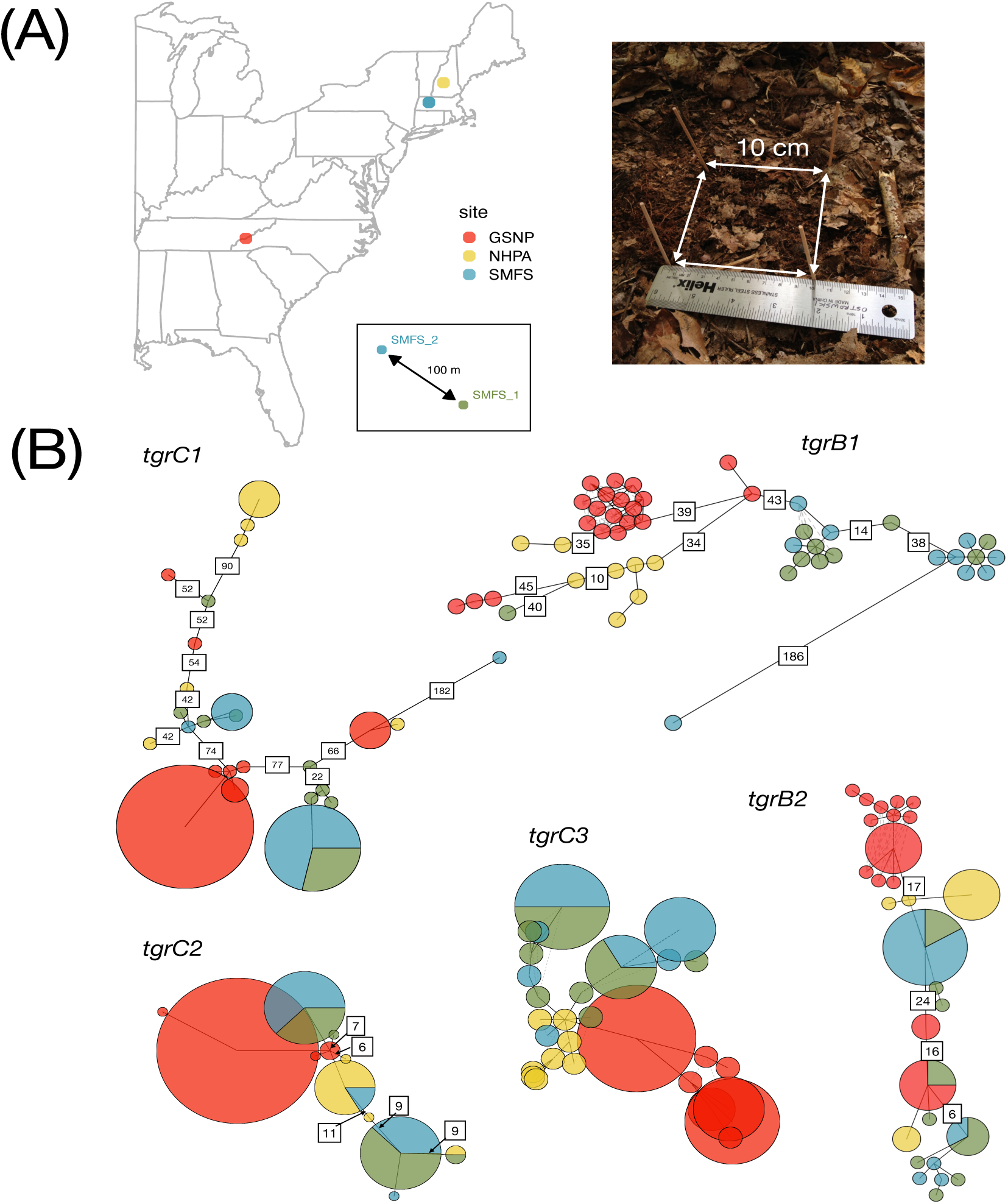
Centimeter-scale spatial variation in *tgrBC* genes. **(A)** Sampling locations for 48 *D. discoideum* strains, GSNP: Great Smoky National Park; NHPA: New Hampshire, Proctor Academy; SMFS: Smith MacLeish Field Station. Each site consisted of a 10-cm by 10-cm plot, from which surface soil was collected for isolation of *Dictyostelium*. **(B)** Haplotype networks for *tgrB* and *tgrC* genes, illustrating the centimeter-scale diversity. Each circle represents a unique haplotype, and its area is proportional to the number of strains that have that haplotype. Edges and numbers indicate the inferred mutational steps separating the two haplotypes; for simplicity, numbers are only shown for edges with distances >4. For highly polymorphic genes *tgrB1*, *tgrB2*, and *tgrC1*, each 10-cm by 10-cm

At each site, we collected 6 g of soil from the surface of a 10-cm by 10-cm plot. Thus, the maximum distance between any pair of isolates from the same soil sample is 14 cm (= √(10^2^ + 10^2^)). Our prior sequencing, including strains from these locations, indicated low genetic differentiation at distances of tens to hundreds of meters and very high differentiation beyond one kilometer (mean *F*_ST_>0.4; Kuzdzal-Fick, in preparation). Thus, we expect some gene flow between SMFS1 and SMFS2, but little gene flow among the more distant pairs of sites.

For nucleotide diversity, our findings with the local strain collections mirrored what we observed with the geographically widespread Nanopore genomes: high polymorphism (in descending order) in *tgrC1*, *tgrB2*, and *tgrB1*—and substantially lower polymorphism in *tgrC2*, *tgrC3*, and *tgrC5* (Fig. 8). Put another way, the local *tgrC1* and *tgrB2* polymorphism is roughly five times higher than the genome average based on geographically widespread strains (e.g., 0.015/0.003 = 5). We observed similar patterns of polymorphism among across populations, suggesting that selection is operating similarly across geographically independent sites. Especially notable was the consistently high polymorphism in *tgrB2*, which is not known to have a role in kin recognition.

**Figure 8.**
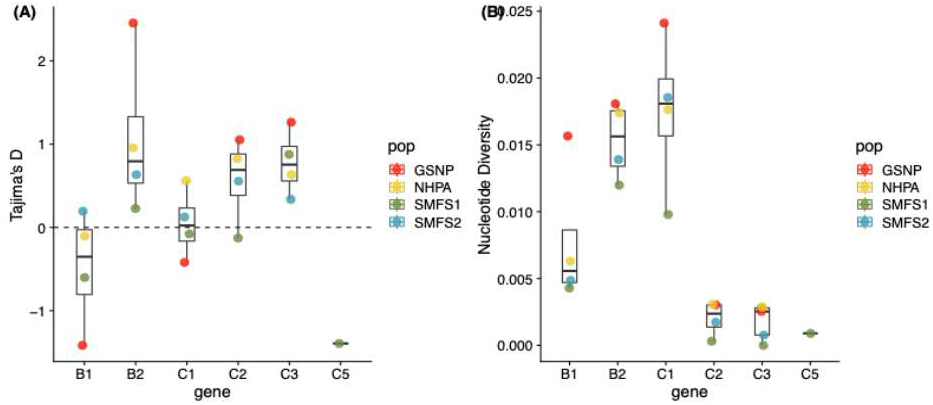
Molecular evolution analyses of *tgrB* and *tgrC* genes. (A) Tajima’s *D*, a metric of selection that typically ranges from approximately +2 to -2 but can be more extreme. Values of zero are expected for genes that are evolving neutrally, i.e., in the absence of selection. Strongly positive values, particularly compared to other genes in the genome, can indicate balancing selection – i.e., an excess of intermediate-frequency alleles. Strongly negative values, compared to other genes in the genome, can indicate positive or purifying selection –.e., skewed allele frequencies, where a few variants are high-frequency, but most variants are rare. Prior work indicated that the mean of the genomewide distribution of Tajima’s D in *D. discoideum* is negative (Ostrowski et al. 2015), consistent with population expansion. In the current study, the mean value for the genome assemblies from this study is also negative (*D* = -0.25; 95% confidence interval = [-1.56, 1.65]). (B) Nucleotide diversity (π) within local populations. Each point is the mean of eight to twenty natural isolates per site sequenced using the Sanger technology.

To further examine how selection is operating on these genes, we calculated Tajima’s *D* for each locus (Fig. 8). Large negative values of Tajima’s *D* indicate highly skewed allele frequencies, potentially generated by positive or purifying selection. Conversely, large positive values of Tajima’s *D* indicate multiple alleles at intermediate frequency, potentially generated by balancing selection (Oleksyk et al. 2010; Vitti et al. 2013). Importantly, Crozier’s Paradox predicts that recognition genes will show positive frequency-dependent selection, reflected in depleted genetic variation and a negative Tajima’s *D*. Conversely, effective maintenance of genetic variation should be reflected in positive values of Tajima’s *D*.

Despite the large differences in levels of polymorphism between *tgrB2*, *tgrC2*, and *tgrC3*, they all showed similarly positive values of Tajima’s *D*. However, *tgrC1* has a Tajima’s *D* value close to zero, a surprising result given its extreme polymorphism thought to be driven by balancing or diversifying selection. Furthermore, *tgrB1* showed a consistently negative Tajima’s *D*, potentially indicating the variation-removing selection predicted by Crozier’s Paradox. Overall, Tajima’s *D* analysis suggests differences in how selection is operating on these genes in nature that were not evident from polymorphism analyses alone.

To visualize the spatial distribution of genetic variation in the *tgrB* and *tgrC* genes, we created haplotype networks (Fig. 7B). These networks for *tgrB1*, *tgrB2*, and *tgrC1* reveal many haplotypes separated by long branches, indicating highly diverged alleles. Importantly, each 10-cm by 10-cm plot harbored multiple genetically diverged haplotypes differing by more than a hundred nucleotides. Haplotypes from the same site (same color) are often not directly connected to each other by an edge. These results indicate that *tgrB* and *tgrC* genes are not only extraordinarily polymorphic across the species as a whole, but a high fraction (∼38%) of the species-wide polymorphism exists in just a few grams of soil within each locality.

Different gene regions—particularly different protein domains—can vary in their selective constraints. In addition, in the case of Tajima’s *D*, averaging positive and negative values (resulting in a value of zero) could lead to erroneous conclusions of no selection. For that reason, we calculated nucleotide diversity (π), Tajima’s *D*, and *F*_ST_ in sliding windows across the gene, using 100-bp windows and a 25-bp slide (Fig. S6) for the Sanger sequences from the four sites. These results highlight several features of the genetic variation: (1) *tgrC1*’s polymorphism peaks in its 5’ immunoglobulin domains, (2) although *tgrB1* does have a consistently negative Tajima’s *D* across most of the gene, the region surrounding its transmembrane and cytoplasmic signaling domains shows positive values, indicating probable balancing selection in that specific region, (3) although polymorphism is very low in *tgrC2* and *tgrC3*, they have extremely high values of *F*_ST_ (=1) over some portions of the gene, mostly overlapping the immunoglobulin domains, indicating strong geographic divergence and potentially local adaptation. These results indicate diverse patterns of molecular evolution for the genes despite their close proximity on the chromosome.

The molecular evolution analyses also revealed that much of *tgrB2*’s extreme nucleotide diversity—which reaches a peak higher than any other *tgrBC* gene and twice as high as the maximum reached by *tgrB1* and *tgrC1*—can be attributed to a short stretch of just ∼100 bp near the start of the gene (Fig. S7). Sequence variation in this region segregates largely into a couple of highly divergent haplotypes at high frequency. The region of extreme divergence is discrete and surrounded by regions of high sequence homogeneity. The resulting mosaic appearance of *tgrB2* suggests genetic variation that has been introduced suddenly rather than accumulated slowly through single mutations. However, sequence comparisons did not identify a clear source, as the region was equally similar to *tgrB1* and *tgrB2*. Phylogenetic analyses of the *tgrB1*, *tgrB2*, and *tgrB_p*s, however, showed intermingled paralogs within clades, a signature of interlocus gene conversion or recombination (Fig. S7).

To assess whether *tgrBC* sequence variation is generated through recombination or gene conversion, we used GARD to detect intralocus recombination and identify potential breakpoint locations. These analyses support *intralocus* recombination for *tgrB1* and *tgrC1* (cAIC*_B1_*=391_;_ cAIC*_C1_*=12039, respectively), based on the significantly better fit of models that allow phylogenetic incongruence, but not for *tgrB2*. The best estimate of breakpoints locations occurred around position 806 in *tgrB1* (within an Ig domain) and position 1290 in *tgrC1* (between two Ig domains; Fig. S8).

GARD operates only on single-locus alignments, so it detects only intralocus recombination. To explore potential interlocus exchange, which occurs frequently among tandemly duplicated genes (Dumont 2015; Harpak et al. 2017), we used several different methodologies implemented in openRDP, which vary in their sensitivity to older versus more recent events. These results also indicate both intra- and inter-locus recombination events, although they disagree on their locations (Fig. S8). In addition, 3Seq identified a region of interlocus recombination in *tgrB2*, which coincides well with the highly diverged alleles region of extreme (*π*=0.07), supporting our hypothesis that the highly discrete mosaicism observed in this gene has been generated by recombination among paralogs.

### Dynamic evolution and mosaicism in the *tgrBC* locus

Dynamic evolution of the gene cluster is apparent among our strains, with the most noticeable differences between the *D. discoideum* and outgroup strains (EO1176 and EO1089; Fig. 9). Nanopore assembly, as well as Sniffles structural variation analysis, also confirmed the suspected deletion of *tgrB2* from its expected genomic location (Fig. 9). Liftoff analyses of the QS18 and QS33 assemblies suggested that *tgrB2* was present but in a different location: the locus where *tgrB_ps* resides in AX4 and other strains, making it paired with *tgrC_ps1*. Examination of the sequence surrounding the translocated *tgrB2* alleles revealed about 50 bases at the end of the gene that do not match the non-translocated *tgrB2* alleles, suggesting a cut-and-paste event. These non-matching bases are nearly identical to a stretch of *tgrB1*, suggesting that the *tgrB2* alleles acquired portions of *tgrB1* through the transposition event. However, beyond those bases, the *tgrB2* bears no further resemblance to *tgrB1*.

**Figure 9.**
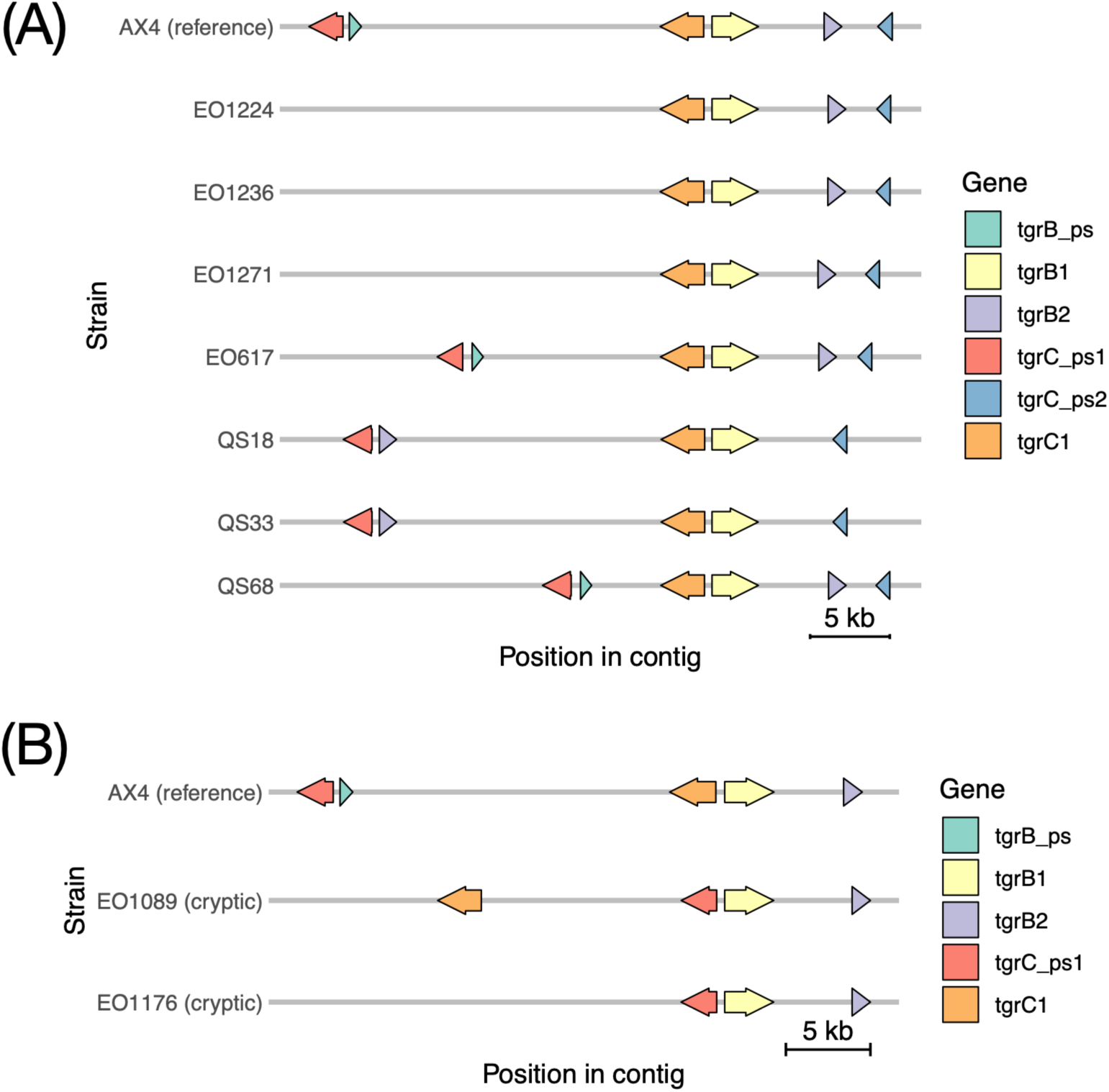
Dynamic gene family evolution in the *tgrBC* locus. Each line shows the inferred location of the *tgrBC* genes and their chromosomal locations, based on Liftoff analysis and centered on *tgrB1* and *tgrC1*; intervening non-*tgrBC* genes not shown. Genes are named according to their match in the reference strain AX4. **(A)** *D. discoideum* strains (and one cryptic species strain) compared to the canonical AX4 arrangement. QS18 and QS33 show a translocation of *tgrB2*, consistent with our earlier PCR results showing a deletion impacting this gene (see Fig. S3). **(B)** Section of the *tgrBC* locus, comparing the cryptic species to the canonical AX4 arrangement.

The molecular evolution analyses, indicating high polymorphism in *tgrB2*, suggests that it might have a role in kin recognition. Transcriptomic analysis of QS33, which harbors the non-syntenic *tgrB2* also reveals that: (1) the allele is as strongly expressed, at the same time in development as *tgrB1* and *tgrC1* (<u>Table S4</u>), (2) the non-syntenic *tgrB2*, like the syntenic *tgrB2*s in other strains, is a truncated *tgrB* protein lacking a predicted transmembrane domain, and (3) despite potentially having a paired *tgrC* (which the syntenic *tgrB2’s* lack; compare locations of *tgrB2* in Fig. 9A), the neighboring *tgrC* gene appears to be a pseudogene, just as it is in other strains. Specifically, although the *tgrC* mRNA is spliced in QS33, there are internal stop codons present and the expression level was low compared to *tgrB1*, *tgrC1*, and *tgrB2* (<u>Table S4</u>).

At first glance, the cryptic outgroup strains, EO1089 and EO1176 appear to lack a functional *tgrC1* that is paired with *tgrB1* — instead, the *tgrC* paired with *tgrB1* is labeled as *tgrC_ps1*, a pseudogene in AX4. Further inspection of these alleles, however, reveals that the EO1089 allele is a full-length *tgrC* gene. Using BLAST, we also found two unpaired *tgrC* genes upstream of the *tgrB*-*tgrC* pair (Fig. S9). We examined these three copies, and they lack stop codons once spliced. They differ from one another in sequence and have indels that are all in multiples of three. Thus, EO1089 has a single *tgrB*-*tgrC* head-to-head pair, but multiple upstream copies of *tgrC* (Fig. S9).

The EO1176’s sole *tgrC* allele was similar to that of EO1089 in sequence; the two alleles differ at only 28 nucleotides across 2674 base pairs, consistent with the strains’ close phylogenetic relationship (Fig. 3). However, the EO1176’s *tgrC* allele has five apparent frameshift mutations compared to the in-frame EO1089 allele. We noticed that these putative indels occurred in short homopolymers, tandem repeats, or other low-complexity sequence—regions that are prone to sequence errors. We therefore examined the underlying reads and found that none of the indels are strongly supported (Fig. S10). In four of five cases, the reads best support the EO1089 allele (i.e., no indel). In the fifth case, the reads indicate a potential deletion relative to the EO1089 allele, but the deletion would maintain the reading frame. Thus, with EO1176 as the only possible exception (and poor support for it being an exception), every strain harbors multiple *tgrB* and *tgrC* full-length genes as well as gene fragments, but only a single, full-length *tgrB*-*tgrC* head-to-head pair.

## Discussion

### Social amoebae show extraordinary levels of centimeter-scale sequence diversity in allorecognition loci—indicating the means and opportunity for selection on kin recognition in nature

Evolutionary biologists have long noted the potential for conflict in the *Dictyostelium* life cycle, given that its transition to multicellularity involves a stage where a fraction of the population dies altruistically to support the rest (Strassmann et al. 2000; Strassmann and Queller 2011; Ostrowski 2019). The ability to recognize and evade costly cooperation with unrelated strains through interactions at the *tgrB1*-*C1* locus is thought to be a valuable protection against cheating (Ho et al. 2013; Ågren et al. 2019; Ostrowski 2019). Nevertheless, despite high levels of species-wide polymorphism in these loci (Benabentos et al. 2009), it has remained unclear if sufficient variation is present over the small spatial scales where the decision to cooperate or reject must take place.

Our results reveal high levels of sequence variation in allorecognition at the centimeter-scale, over distances traversable by the migratory slugs and within the known range of genetic migration (Jack et al. 2011; Jack et al. 2015). Simply put, kin recognition polymorphism must occur at specific spatial scales, where strains encounter individuals of varied (i.e., both high and low) relatedness, making the ability to distinguish between them selectively important. This crucial requirement has been met here, with extremely divergent (as well as identical) alleles co-occurring over distances of not more than 14 cm in the soil.

### Is selection depleting variation through positive-frequency dependence, as suggested by Crozier’s Paradox?

Crozier’s Paradox predicts that genetic kin recognition will be a victim of its own success, depleting its own genetic variation through positive frequency-dependent selection (Crozier 1986; Rousset and Roze 2007). Here, we show that neither *tgrB1* nor *tgrC1* is depleted of genetic variation. Indeed, both genes exhibit high levels of sequence variability, with nucleotide diversity values in the top 0.3% of the genome-wide values. While genome-wide nucleotide diversity, based on geographically distant strains, provides an estimate of 4% for these genes, isolates from the single soil samples have values around 1.5-2%. Given that many of these pairwise comparisons will be zero (for co-occurring isolates that arise by recent cell divisions), a large fraction of species-wide allele divergence can be found at the centimeter-scale in nature. Our findings also hold across geographically distant sites, indicating consistent patterns of genetic variation and selection in nature.

Our results do not show strong evidence of variation-depleting selection. Tajima’s *D* values tended to be positive for *tgrB2*, *tgrC1* and the other *tgrC* genes. Noting that the mean Tajima’s *D* is negative in this and our previous study (Ostrowski et al. 2015), this pattern is more consistent with balancing selection. However, Tajima’s *D* values were negative across most of *tgrB1*. It showed high dN/dS near the start of the gene (Fig. 6), where the polymorphism is greatest (<u>Fig. S6</u>). Combined with lower polymorphism compared to *tgrC1* and *tgrB2* (albeit still very high overall), these results suggest that *tgrB1* may not maintain its genetic variation as well as these two other genes. The dN/dS analysis suggests the reason: purifying selection on its 3ʹ end. *tgrB1* differs from both *tgrC1* and *tgrB2* in having a cytoplasmic domain at the end of the gene, used for intracellular signaling. This domain ensures information from the externally bound TgrC1 is transmitted internally to enable responsive cellular decision-making (Katoh-Kurasawa et al. 2024). Our results thus suggest a selective conflict between the two parts of this bifunctional gene. While this conflict may constrain its evolution, it does not appear to be caused by Crozier’s Paradox itself, which predicts loss of genetic variation through positive frequency-dependent selection, specifically. Overall, we do not see strong evidence of the existence of the paradox, in the form of positive frequency-dependent selection, or the outcome of the paradox, in the form of depleted genetic variation.

### Molecular evolution analyses uncover additional targets of selection, especially *tgrB2*

Although the established kin recognition genes *tgrB1* and *tgrC1* show extraordinary sequence variability, our analyses uncovered additional genes and members of the *tgr* gene family that are highly polymorphic or show other signatures of selection. *tgrB2* is especially perplexing: its sequence variation is similar in magnitude to that of *tgrB1* and *tgrC1* overall, and its peak of polymorphism near the N-terminus dwarfs that of the other two genes. The high frequency of its haplotypes, especially across multiple populations, suggests they are unlikely to be selectively neutral. While not strongly expressed in the standard lab strain, AX4 (Fig. S2), the *tgrB2* gene is strongly expressed during development in QS33 (<u>Table S4</u>). Its strong expression, at the same time during development as *tgrB1* and *tgrC1*, demonstrates that we cannot yet rule out a functional role for other *tgr* genes, including in kin recognition itself.

### Variable selection across the *tgrBC* locus and recombination

Holland and colleagues (2025), using *D. discoideum*’s closest outgroup species, have shown that the *tgrBC* locus is hypermutable, with duplications, deletions, and rearrangements that rapidly generate novel alleles and loci, helping to explain how these genes generate their extreme sequence polymorphism. With highly related and physically co-located strains, we likewise show evidence of rearrangements and a highly dynamic genomic locus.

Against this backdrop of an unstable genomic region, however, we find variation among the different *tgrB* and *tgrC* genes in their likely form of selection and levels of genetic variation. These results strongly suggest that the hypermutability is nevertheless shaped by natural selection, with *tgrB1*, *tgrC1*, and *tgrB2* under stronger selection than the rest of the genome or even other *tgr* genes in the same locus (*tgrC2*, *tgrC3*). Locally restricted recombination (see Holland et al. (2025)), may explain how tightly linked genes, especially *tgrB1* and *tgrC1*, can evolve independently. The high rate of recombination enables rapid molecular evolution but can also confine it to particular genes—or even gene regions, as in *tgrB1*—where its recombination breakpoint separates the two sections of the gene experiencing divergent types of selection (positive vs purifying).

### Gene family dynamics can replenish lost genetic variation

Prior theory indicated that typical mutation rates are too low to compensate for the speed of genetic variation loss at kin recognition loci (Rousset and Roze 2007). Crucially, this work did not consider the elevated rates of variation that arise in large gene families through gene duplication, recombination, and gene conversion. Gene conversion and non-allelic recombination can generate novel sequences and increase haplotype diversity, particularly in gene families under strong diversifying selection (Spurgin et al. 2011; Klitz et al. 2012; Daugherty and Zanders 2019; Minias 2024). Thus, even if only one *tgrB–tgrC* pair mediates recognition at any given time, duplicate genes can create a reservoir of variation, as well as an opportunity for recognition to shift to a novel locus if it becomes a more reliable indicator of relatedness (Scott et al. 2023). While the dynamics of multigene families are well-recognized in the evolution of immune recognition, to our knowledge, they have not previously been proposed as a solution to Crozier’s Paradox.

In conclusion, our results indicate high genetic variation in *Dictyostelium*’s kin recognition genes at the relevant spatial scales in the soil, suggesting that this organism evades Crozier’s Paradox. Our results provide further support that genetic kin recognition is a mechanism that can stabilize altruistic cooperation.

## Methods

### Soil Sampling and Isolation of EO Strains

– EO strains were isolated from soil samples obtained from the top centimeter of soil in 10-cm by 10-cm plots in the following locations: Proctor Academy, New Hampshire (NHPA); Smith-MacLeish Field Station (SMFS1, SMFS2), and Great Smoky National Park (GSNP). GPS coordinates are provided in <u>Table S1</u>. To isolate strains, we combined 6 g of soil with 30 ml of sterile water and then deposited approximately 200-400 ul across multiple hay infusion agar plates <u>(Cavender and Raper 1965)</u> along with 400 µl of stationary phase culture of *Klebsiella pneumoniae* grown in SM broth. Once slugs and fruiting bodies were observed, we collected spores from a single fruiting body per plate, deposited each one in separate wells of a multi-well plate, containing KK2 buffer and 30% glycerol, and froze the plate at -80C.

### Cloning of EO Strains

– Naturally formed fruiting bodies can potentially contain more than one strain. To obtain clonal cultures, we plated spores from the freezer stock at low density on lawns of *K. pneumoniae* on SM agar plates, so they form plaques, i.e., clearings in the bacterial lawn. For each isolate, we collected fruiting bodies from the center of a well-isolated plaque and transferred the spores to a fresh SM plate with *K. pneumoniae* to expand the population. We collected the spores and froze them at -80C in KK2 buffer with 30% glycerol.

### Cryptic outgroup strains

– Two of these strains, EO1089 and EO1176, were identified using previous Illumina sequencing as likely cryptic species (Kuzdzal-Fick et al., in preparation). Briefly, these strains are highly genetically divergent from our *D. discoideum* strains (Fig. 3), but more closely related to *D. discoideum* than *D. discoideum* is to its known sister species, *D. citrinum*. These outgroup strains co-occurred in soil samples with *D. discoideum*. They also lacked any discernable morphological differences from *D. discoideum:* they formed stalkless slugs of the expected size and well-isolated, yellow fruiting bodies that were upright with a basal disk.

### QS strains

– QS strains were collected previously by the Queller-Strassmann lab and sequenced initially by Ostrowski et al. (2015) using Ilumina and Benabentos et al. (2009) using Sanger sequencing. We sequenced some of these strains again using Sanger, Illumina, and/or Nanopore sequencing. Further details are provided in <u>Table S1</u>.

### Sanger Sequencing and Analyses

– We sequenced 48 strains at *tgrB1*, *tgrB2*, *tgrC1*, *tgrC2*, *tgrC3*, and *tgrC5* using Sanger sequencing (<u>Table S1</u>), obtained from 10-cm by 10-cm plots at four sites (Fig. 7A). Detailed methods for PCR amplification and sequencing are described in the supplement. We generated consensus sequences for each strain and aligned them using the MUSCLE algorithm in Geneious 11.1.4 (Edgar 2004). For coding sequences, we spliced out the introns based on the AX4 gene model. Analyses of nucleotide diversity, Tajima’s *D*, and *F*_ST_ were carried out using PopGenome v.2.7.7 in R (Pfeifer et al. 2014). FASTA alignments were transformed into overlapping sliding windows of 100 bp, with 25-bp steps. We used the “neutrality.stats” and “diversity.stats” commands to obtain the values for Tajima’s *D*, *F*_ST_, and nucleotide diversity for each gene and sliding window. Haplotype networks were made using the R package PEGAS v. 1.4 (Paradis 2010). We manually adjusted the placement of the edges to prevent overplotting of the nodes.

### Nanopore Sequencing

– Genomic DNA was isolated according to the protocol in Ostrowski et al. (2015) and stored in TE buffer (pH 8.0) at 4°C until sequencing. Sequencing was performed on Oxford Nanopore Technologies (ONT) MinION Mk1B devices using FLO-MIN106 (R9.4.1) flow cells. Both new and reused cells were used, with reused cells washed between runs using the ONT Flow Cell Wash Kit (EXP-WSH004). Libraries were prepared from ∼1 µg gDNA with the ONT Genomic DNA by Ligation Protocol (SQK-LSK109, version 14Aug2019) with the following modifications. Prior to library preparation, DNA fragments shorter than 10 kb were removed with a Short Read Eliminator (SRX) kit (Circulomics). During AMPure bead clean-up steps, incubation on the magnetic rack was extended to 2–4 min until the liquid was clear. For the final elution, the library was diluted in buffer pre-heated to 50 °C and incubated for 10 min.

### Nanopore Genome Assembly

– Reads for all strains were basecalled using Guppy (v. 6.3.8), with a minimum quality score filter of 9. Draft assemblies were generated *de novo* with Flye v2.9 (Kolmogorov et al. 2019) without specifying a target genome size. The resulting assemblies were polished four times with Racon v1.5.0 (Vaser et al. 2017), followed by a single polishing step with Medaka (v1.7.2; https://github.com/nanoporetech/medaka). For all genomes excluding the two cryptic strains, a circular *Klebsiella pneumoniae* contig (∼5.2Mbp) was generated, which was removed manually before calculating genome statistics.

Polished assemblies were initially annotated by lifting over annotations from the AX4 reference genome available from DictyBase (http://dictybase.org/) using Liftoff v1.6.3 (Shumate and Salzberg 2021), with multicopy mode enabled. Annotated assemblies were scaffolded to the AX4 reference genome with RagTag v2.1.0 (Alonge et al. 2022). QUAST v5.3.0 (Mikheenko et al. 2023) was used to generate assembly statistics on both scaffolded and unscaffolded assemblies. BUSCO v5.5.0 (Simão et al. 2015) was used to assess the genome completeness of all sequenced strains. All BUSCO assessments were run with the ‘--auto-lineage-euk’ flag to automatically select the best database for comparison, which selected the eukaryote_odb10 database in all cases.

To assess tgr gene content in the *tgrBC* cluster, we used tblastn (BLAST+ 2.13.0) with AX4 *tgr* proteins as queries against nucleotide databases built from each strain’s contig-level assembly (-evalue 1e-5, -max_target_seqs 500, -seg yes). High scoring pairs (HSPs) from the same query were merged into alignment blocks, requiring collinearity in both query and subject coordinates with a maximum subject gap of 3 x (query gap) + 300 bp to allow for introns. Overlapping alignment blocks were then merged into loci. Each locus was assigned to a gene family (e.g., *tgrB* vs *tgrC*) based on a similarity metric, calculated as min(qcov, lcov)*pid, where qcov and lcov are the union of interval coverages of the query and locus, respectively, and pid is the length-weighted mean percent identity across HSPs. Because tblastn coordinates are based on contig assemblies but Liftoff coordinates are based on RagTag scaffolds, we compared them by anchoring on the tgrB1 locus.

We used a custom MAKER pipeline and reciprocal best hits to identify orthologous genes. Alignments were generated using MAFFT v7.526. Alignments of orthologs in all *D.discoideum* strains were used to calculate nucleotide diversity (π) for all genes across the genome with Egglib v3.3.5 (Siol et al. 2022) and only sites with less than 30% missing data were retained. dN/dS was calculated using PAML v4.10.7 (Yang 2007), and McDonald-Kreitman tests were carried out using mkado v. 0.1.0 (Rivera-Colón et al. 2026). Further details of analyses are provided in the supplemental methods.

## Data and software availability

Assemblies and Sanger sequences have been uploaded to NCBI under BioProject PRJNA1446954. Pipelines, scripts, and datafiles are available on Github (https://github.com/eostrow/tgr-genes) and will be archived on Zenodo prior to publication (DOI: TBD).

## Acknowledgements

The authors would like to acknowledge the use of the Research Computing Data Core (RCDC) at the University of Houston and the New Zealand eScience Infrastructure (NeSI) high-performance computing facilities for data analyses. New Zealand’s national facilities, provided through NeSI, are funded jointly by NeSI’s collaborator institutions and through the Ministry of Business, Innovation & Employment’s Research Infrastructure program. EAO and MC were supported by a grant MFP-UOA2412 from the Marsden Fund Council, managed by the New Zealand Royal Society Te Apārangi. CB and SM were supported by Massey University Doctoral and Masters scholarships, respectively. TCL and MH were supported by National Institutes of Health grant R35GM150681. The authors would like to thank Proctor Academy, Smith MacLeish Field Station, and Mountain Lake Biological Station for permission to collect soil samples. Soil from Great Smoky National Park was collected under permit GRSM-2017-SCI-2014.

