## Supplement for "Evolutionary dynamics of the *tgr* gene family in *Dictyostelium* allows escape from Crozier’s Paradox"

[*] Contributed equally

### Supplemental Methods and Approach

***Sequencing Strategy*** - Our sequencing strategy was informed by initial observations that QS strain *tgr* gene sequences generated with Illumina (Ostrowski et al. 2015) did not match those generated by Sanger sequencing (Benabentos et al. 2009). Manual inspection of the Illumina sequences indicated read mismapping across paralogs caused by low sequence divergence among paralogs and high allelic diversity with paralogs, which could not be remedied with more stringent filters. We initially focused on resequencing with Sanger, which generated single bands, high-quality chromatograms, and repeatable sequences, indicating high accuracy. Long-read sequencing offered an ability to obtain *de novo* genome assemblies that completely circumvented read mapping; therefore, Sanger and Nanopore represent orthogonal approaches to obtaining accurate *tgr* gene sequences, with the latter presenting an opportunity to detect previously unrecognised paralogs and structural rearrangements that are common in this type of gene family.

We wanted to remedy several limitations of earlier work by us and others, which can make detecting selection challenging (Ostrowski et al. 2015; Noh et al. 2018; de Oliveira et al. 2019). First, *D. discoideum* has extremely low polymorphism, which means that a large fraction of genes have few or no SNPs, and dN/dS is often undefined because dS=0. The closest known outgroup, *D. citrinum*, is highly diverged, had a fragmented assembly, and lacked many *D. discoideum* orthologs, creating an intrinsic bias in which genes could be compared. Second, molecular evolution statistics that make use of the allele frequency distribution or differences (e.g., Tajima's $D$, $F_{ST}$) can be powerful for these closely related strains that have few distinguishing SNPs. However, existing strain collections were not suitable for estimating allele frequencies, given uncertain spatial relationships among the strains only and one or a few isolates were available per site. For this reason, we undertook *D. discoideum* isolation with our ideal tests of molecular evolution in mind: we sampled numerous isolates over the small spatial scales where gene flow occurs (14 cm maximum distance), as well as large spatial scales (~1,000 km apart) to capture independently evolving populations.

Illumina genome sequencing of our large (>700+) natural isolate collection also uncovered a distinct genetic cluster that was highly divergent from the vast majority of our isolates, but also much more closely related to them than their closest known outgroup, *D. citrinum*. We thus opted to sequence and assemble two of these strains, with the goal of obtaining a new outgroup sequence that would permit tests of selection acting over longer timescales (e.g., dN/dS). In summary, our final sequencing dataset comprises *tgrB1*, *tgrB2*, *tgrC1*, *tgrC2, tgrC3*, *tgrC5* in 48 EO strains from four soil samples at three sites and nine *de novo* assemblies, including two novel outgroup strains.

### Supplemental Methods

#### Sanger sequencing

***Strains, culture conditions, and DNA extraction*** – Spores were inoculated from frozen spore stocks onto SM agar plates in co-culture with the *K. pneumoniae.* Cells were allowed to develop into multicellular fruiting bodies. DNA was extracted by placing 5-10 sori in 250 μL of a 5% Chelex solution, followed by the addition of 10 μL of 10 mg/ml Proteinase K. The samples were then mixed and incubated at 40°C for 4 h, followed by 70 °C for 10 min to denature the proteinase K. The supernatant containing the genomic DNA was stored at -20 °C until use.

***Primer design and PCR protocol*** – The genes (*tgrB1, tgrB2, tgrBps, tgrC1, tgrC2, tgrC3,* and *tgrC5*) were each amplified using a single pair of primers for each gene, from genomic DNA. Primers were designed in Primer3 based on the AX4 reference genome (Eichinger et al. 2005). The PCR protocol consisted of an initial denaturation step at 94°C for 120 s, followed by 35 cycles of: a denaturation step at 94°C for 20 s, two annealing steps for 15 s each (initial 5°C lower than primer melting temperature ($T_m$) and secondary 8°C lower than primer $T_m$), and an extension step at 62 °C for a variable time depending on the length of the gene (extension time = 1 min/Kb * 1.30). Gel electrophoresis was used to visualize and verify the presence of a single band of the expected size, using a 2% agarose gel, stained with 0.5 µg/mL of ethidium bromide. Additional sequencing primers were designed so that the entire gene would be covered by at least one read.

***PCR product purification and DNA sequencing*** – PCR products were purified using a QIAGEN PCR Purification Kit (protocol according to manufacturer's specifications and eluted in water) with QIAGEN purification columns. Some products were instead purified directly by GENEWIZ Inc., which used enzymatic PCR purification (ExoSAP-IT). Purified PCR products were sent to GENEWIZ for Sanger sequencing.

#### Illumina sequencing

Beginning with a multisample vcf file restricted to biallelic alleles (Kuzdzal-Fick et al. 2023), we used bcftools (1.14) to calculate variant allele frequency with +fill-tags -t FORMAT/VAF. After filtering for –minDP 4 and –maxDP 510 with vcftools (0.1.16), we ran gatk (gatk4-4.2.5.0) VariantFiltration --genotype-filter-name "VAF_filter"  \ --genotype-filter-expression "isHomVar == 1 && VAF < 0.8".  We used samtools (1.11) faidx to select each region of interest and bcftools consensus --sample -M 'N' to produce fasta files of each strain's *tgr* sequences.

***Sequence accuracy analysis*** – Sequences obtained through Illumina and Nanopore sequencing were compared to a Sanger counterpart to assess accuracy over the *tgr* gene regions. Comparisons were made by aligning gene sequences with MAFFT v7.526 (Katoh et al. 2002) and counting the mismatched base pairs. Alignments of orthologs in all

*D.discoideum* strains were used to calculate nucleotide diversity (π) for all genes across the genome with Egglib v3.3.5 (Siol et al. 2022) and only sites with less than 30% missing data were retained.

***Non-synonymous to synonymous mutation ratio estimation*** – When estimating non-synonymous to synonymous mutation ratios, an alternate gene annotation pipeline was employed to ensure accurate prediction of coding sequences. We used GeneMarkEP+ (Brůna et al. 2020) to predict protein sequences and provided the reference *D. discoideum* AX4 protein sequences as the protein hint database. To identify rRNA genes, we used RNAmmer (Lagesen et al. 2007) with default parameters. We then used MAKER (Cantarel et al. 2008) to generate a custom repeat library, which was provided to RepeatMasker for softmasking each genome prior to annotation. tRNA genes were identified using tRNAscan-SE (Chan et al. 2021). Functional annotation labels were added to each predicted coding sequence with results from BLASTP of the UniProt database release 2024_01 (The UniProt Consortium 2024) and InterProScan v5.57-90 (Blum et al. 2025) of the PFAM database (Pei et al. 2024). Finally, unique gene identifiers were appended using scripts from the MAKER pipeline.

Orthologs were identified from the MAKER set of coding sequences using mmseq2 v17.b804f (Steinegger and Söding 2017) 'easy-rbh' reciprocal best hits to the reference AX4 coding sequences. Codon-aware alignments between orthologs were produced using MAFFT and pal2nal v14.1 (Suyama et al. 2006). Non-synonymous to synonymous mutation ratios were estimated in a pairwise fashion between an outgroup (EO1089/EO1176) and a *D. discoideum* strain for a single omega value (model 0) using PAML/codonml 4.10.7 (Yang 2007). All possible pairs were considered. Sliding window analysis was performed pairwise in 100 codon windows, with a step of 1 codon and the average dN/dS value over all pairwise comparisons per window plotted. Windows which did not have enough variation to estimate dN/dS (dS < 0.01 or dN/dS = 99) were excluded from the calculation.

***Structural variation analysis*** – Sniffles v2.6.0 (Smolka et al. 2024) was used to identify structural variations between our assemblies and the AX4 reference after mapping base-called reads to the final assemblies with minimap2 v2.22 (Li 2018). Overlaps between structural variations and areas of interest were identified with bedtools v2.31.1 (Quinlan and Hall 2010). Recombination was tested using openRDP v.0.1 (https://github.com/PoonLab/OpenRDP), with option -m threeseq rdp. We ran GARD v 0.2 using Datamonkey (https://www.datamonkey.org/GARD), assuming a beta-gamma distribution of site-to-site variation and two rate classes.

### TgrC1 Splicing

In cases where SNPs occurred in or near the splice site consensus sequence, we verified the splicing. Briefly, strains were revived from frozen stocks on SM agar plates overlaid with *K. pneumoniae* culture. Spores from mature fruiting bodies were deposited onto new SM plates with a fresh *K. pneumoniae* lawn and incubated for 40 h at 22 °C in darkness. Cells were collected during mid-exponential growth phase in KK2 buffer, washed three times (450 g, 3 min, 4 °C), pelleted, and then counted. A total of ~$1\times10^7$ to $6\times10^7$ cells was deposited onto mixed cellulose ester (MCE) filters.

Filters were placed atop a cellulose pad (PALL) in a 6-cm Petri dish, moistened with 1 mL cold PDF. Plates were incubated at 22 °C in darkness in sealed, humidified containers. After 9 hours, cells were scraped off the filter into the inner cap of a 1.5 mL Eppendorf tube using a sterile spatula. We added 1 mL of Trizol and vortexed to lyse the cells. After 5 min incubation, 200 µL chloroform was added, followed by vigorous shaking, a 3 min incubation at room temperature, and centrifugation (12,000 × g, 15 min, 4 °C). The aqueous phase of the mixture was transferred to a new tube, mixed with 500 μL isopropanol, incubated for 10 min, and centrifuged again (12,000 × g, 15 min, 4 °C) to pellet RNA. After washing with 1 mL 75% ethanol, samples were stored at –80 °C until further processing.

For *tgrC1* splicing, the RNA was treated with DNAse using the TURBO DNAse kit (Invitrogen) and cDNA was generated using the iScript cDNA synthesis kit. We used a *tgrC1* gene-specific reverse primer that binds at position 2204 of the gene (5' TGGTCCTGAACGAACTCC-3'). The resulting cDNA was used as template to amplify the region surrounding the intron, using forward and reverse primers that bind at bases 55 and 882 of the gene, respectively (5'-TCAATGAATCCTCCAACACCA-3', 5'-ACATTGGACCGGGATCAACTC-3'). The PCR product was sent for Sanger sequencing at Auckland Genomics. The sequences were mapped to the AX4 reference sequence using Geneious Prime to determine splicing.

### Nanopore Direct RNA Sequencing

For direct RNA sequencing, we used RNA isolated from QS33 as described above, after 9 hours of development, corresponding to the expected timeframe for *tgrB1* and *tgrC1* peak expression (see Fig. S2). We used the Oxford Nanopore direct RNA sequencing kit and followed the manufacturer's protocol (SQK-RNA004), using the polyadenylation mode. We obtained ~40 million reads. The reads were mapped to the QS33 genome assembly using the nanoseq pipeline (https://nf-co.re/nanoseq/3.1.0/ (Ewels et al. 2020), and BAMs were visualized in IGV v2.19.5 (Robinson et al. 2011). We used Liftoff (Shumate and Salzberg 2021) to transfer annotations between the AX4 reference genome and the QS33 assembly. For genes of interest, we extracted the reads and assembled them in Geneious to check splicing and reading frame.

**Table S1. Strains, sampling locations, and sequencing technologies.** Asterisks indicate strains that were sequenced by Benabentos et al. (2009). QS strains were sequenced using Illumina by Ostrowski et al. (2015). All other strains were sequenced in the current study. Bold and underlining indicates strains also sequenced using Nanopore.

| Collecting Location | GPS Coordinates (Lat, Lon) | Strains |
|---|---|---|
| Great Smoky National Park, NC (GSNP) | 35.6085, -83.4474 | EO599, EO600, EO601, EO602, EO603, EO604, EO605, EO606, EO607, EO608, EO609, EO610, EO611, EO612, EO613, EO614, EO615, EO616, **EO617**, EO618 |
| Smith McLeish Field Station, Site 1 (SMFS1) | 42.4486, -72.6813 | EO619, EAO620, EO621, EO622, EO623, EO624, EO625, EO626, EO627, EO628 |
| Smith McLeish Field Station, Site 2 (SMFS2) | 42.4491, -72.6821 | EO639, EO640, EO641, EO642, EO643, EO644, EO645, EO646, EO647, EO648 |
| Proctor Academy, New Hampshire (NHPA) | 43.4423, -71.8235 | EO1004, EO1005, EO1006, EO1007, EO1008, EO1009, EO1010, EO1014 |
| Mountain Lake, VA | 37.35, -80.52 | QS1, QS4, QS6*, QS8*, QS9, QS11, QS14, QS15, QS17, **QS18**, QS21*, QS22*, QS23, QS38*, QS45* |
| Houston Arboretum, TX | 29.77, -95.45 | QS31*, **QS68**, QS69, QS70, QS73, QS74, QS80 |
| Little Butts Gap, NC | 35.77, -82.34 | AX4, QS41* |
| Bloomington, IN | 39.22, -86.36 | QS34* |
| Carthage, TX | 32.18, -94.3 | QS30* |
| Effingham, IL | 39.09, -88.58 | QS35 |
| Forrest City, AR | 34.83, -91.47 | QS46* |
| Indian Gap, TN | 35.61, -83.45 | QS39 |
| Land Between Lakes, KY | 39.99, -88.22 | QS36 |
| Linden, TX | 33.06, -94.27 | QS37 |
| Mt. Fuji, Japan | 35.36, 138.73 | QS44* |
| Mt. Greylock, MA | 42.64, -73.17 | QS40* |
| Pasadena, TX | 29.58, -95.07 | QS32* |
| Pasadena, TX | 29.58, -95.07 | QS32* |
| St. Louis, MO | 38.77, -90.18 | QS47* |
| Webster, TX | 29.53, -95.07 | **QS33** |
| Mountain Lake Biol Station, VA (B1) | 37.3536, -80.5362 | **EO1089**, **EO1236** |
| Mountain Lake Biol Station, VA (D2) | 37.3774, -80.5220 | **EO1176** |
| Mountain Lake Biol Station, VA (A1) | 37.3743, -80.5205 | **EO1224** |
| Mountain Lake Biol Station, VA (C1) | 37.3729, -80.5187 | **EO1271** |

**Table S2. Likelihood ratio test results from PAML**

| Gene | Comparison | df | LRT | *P* |
|---|---|---|---|---|
| *tgrB1* | M0 vs M1a | 1 | 272.76 | 2.844e-61 |
| *tgrB1* | M1a vs M2a | 2 | 125.84 | 4.718e-28 |
| *tgrB1* | M7 vs M8 | 2 | 125.47 | 5.67e-28 |
| *tgrB2* | M0 vs M1a | 1 | 18.32 | 1.871e-5 |
| *tgrB2* | M1a vs M2a | 2 | 29.83 | 3.334e-7 |
| *tgrB2* | M7 vs M8 | 2 | 30.04 | 2.996e-7 |
| *tgrC1* | M0 vs M1a | 1 | 211.49 | 6.493e-48 |
| *tgrC1* | M1a vs M2a | 2 | 55.71 | 7.974e-13 |
| *tgrC1* | M7 vs M8 | 2 | 58.73 | 1.763e-13 |

**Table S3. Proportion of sites under selection from PAML**

| Gene | Site class | Proportion | Omega |
|---|---|---|---|
| *tgrB1* | Class 0 (purifying/neutral) | 0.6897 | 0 |
| *tgrB1* | Class 1 (neutral) | 0.2373 | 1 |
| *tgrB1* | Class 2 (positive selection) | 0.073 | 7.9 |
| *tgrB2* | Class 0 (purifying/neutral) | 0.3881 | 0 |
| *tgrB2* | Class 1 (neutral) | 0.6056 | 1 |
| *tgrB2* | Class 2 (positive selection) | **0.0064** | 167.0 |
| *tgrC1* | Class 0 (purifying/neutral) | 0.5597 | 0.06 |
| *tgrC1* | Class 1 (neutral) | 0.3646 | 1 |
| *tgrC1* | Class 2 (positive selection) | 0.0757 | 4.6 |

**Table S4. *Tgr* gene expression in QS33 using direct RNA sequencing.** QS33 cells were collected from filters at 9 hours of development. CPM = counts per million mapped reads. Expression levels were determined using Bambu (Chen et al. 2023) as part of the Nanoseq workflow (https://github.com/nf-core/nanoseq). We also used featureCounts (Liao et al. 2014) to assign reads to features (genes) to accommodate potential transcripts belonging to pseudogenes, which are excluded using standard approaches. Only those *tgr* genes with CPM>0 are shown below. Note that *tgrB2* shows an expression level similar to that *tgrB1* or *tgrC1* (in bold).

| DDBG | Gene | Exonic Length | No. Reads | CPM |
|---|---|---|---|---|
| DDB_G0288711 | tgrM1 | 2562 | 314 | 1990.2 |
| **DDB_G0280531** | **tgrC1** | **2670** | **75** | **475.4** |
| **DDB_G0280693** | **tgrB2** | **990** | **59** | **373.9** |
| DDB_G0277627 | tgrR2 | 2553 | 56 | 354.9 |
| **DDB_G0280689** | **tgrB1** | **2703** | **55** | **348.6** |
| DDB_G0268284 | tgrO4 | 3117 | 38 | 240.8 |
| DDB_G0291668 | tgrA3 | 3267 | 34 | 215.5 |
| DDB_G0268490 | tgrO1 | 2676 | 24 | 152.1 |
| DDB_G0292562 | tgrQ1 | 2016 | 22 | 139.4 |
| DDB_G0291622 | tgrA1 | 2541 | 21 | 133.1 |
| DDB_G0268304 | tgrO3 | 2659 | 21 | 133.1 |
| DDB_G0267654 | tgrO2 | 2670 | 20 | 126.8 |
| DDB_G0286825 | tgrD1 | 2688 | 20 | 126.8 |
| DDB_G0275745 | tgrR1 | 2616 | 15 | 95.1 |
| DDB_G0274225 | tgrA5 | 2805 | 12 | 76.1 |
| DDB_G0284659 | tgrL1 | 2214 | 10 | 63.4 |
| DDB_G0291514 | tgrA4 | 2410 | 10 | 63.4 |
| DDB_G0283335 | tgrI1 | 2568 | 9 | 57.0 |
| DDB_G0292772 | tgrG1 | 2928 | 9 | 57.0 |
| DDB_G0283317 | tgrI2 | 2568 | 7 | 44.4 |
| DDB_G0292732 | tgrF1 | 2754 | 7 | 44.4 |
| DDB_G0281407 | tgrC5 | 2664 | 3 | 19.0 |
| DDB_G0286851 | tgrE1 | 2739 | 3 | 19.0 |
| DDB_G0272736 | tgrN1 | 2835 | 2 | 12.7 |
| DDB_G0283323 | tgrH1 | 3366 | 1 | 6.3 |

**Figure S1. Tgr proteins are named for their distinctive domain structures, which include multiple immunoglobulin folds.** "Tiger" (*tgr*) stands for Transmembrane, IPT/Ig-like E-set Repeat proteins. Shown are the annotations for the TgrB and TgrC proteins. Filled boxes indicate the Uniprot IPT/TIG or Ig E-set domains (IPR002909, IPR014756), whereas shaded blue boxes indicate the homologous superfamily Ig-like folds (IPR013783) based on the Interpro entry.

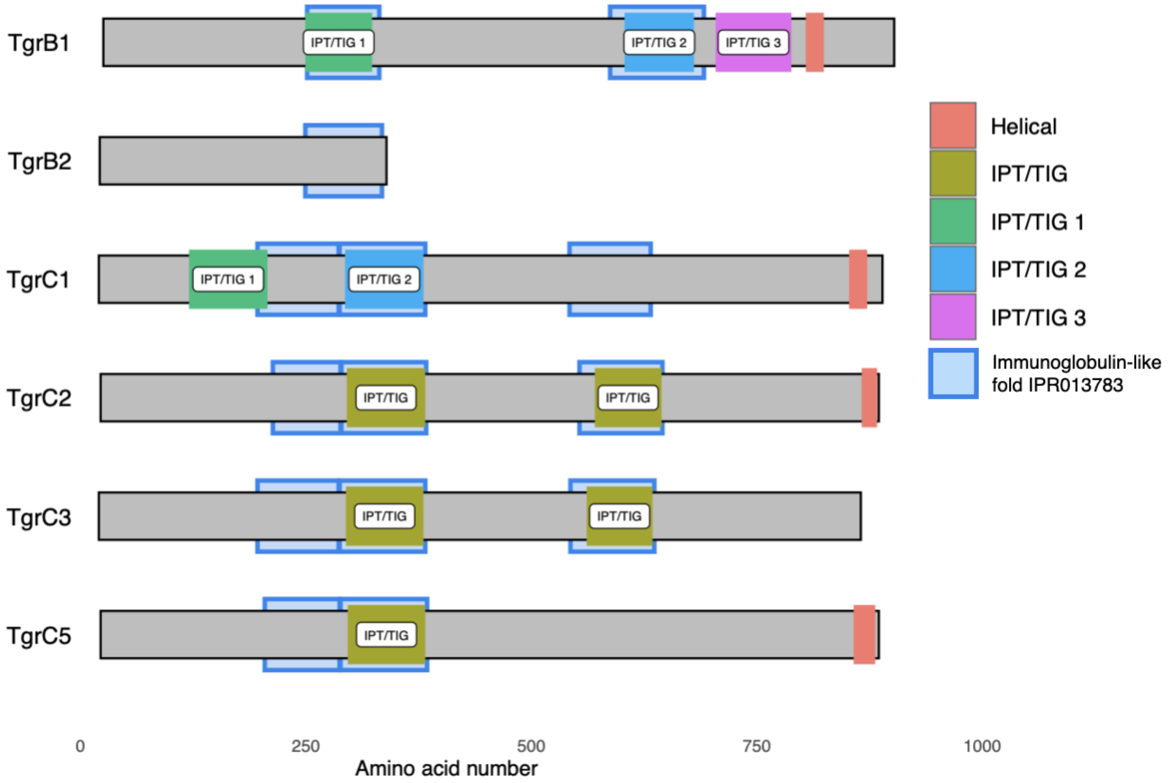

**Figure S2.** Expression of *tgr* genes during clonal multicellular development of the reference (lab) strain AX4, based on Illumina short-read RNA-seq data from Rosengarten et al [(2015)](#) and downloaded from DictyExpress [(Stajdohar et al. 2017)](#). The x-axis is the number of hours since the onset of starvation, with aggregation occurring around 8 hours, slug formation by 12 hours, and fruiting bodies by 24 hours. Y-axis is gene expression, expressed as reads per kilobase per million mapped reads (RPKM). During clonal multicellular development of the lab strain AX4, expression is low or non-detectable for most *tgr* genes except for *tgrB1* and *tgrC1* (the focal kin recognition genes), and *tgrM1*.

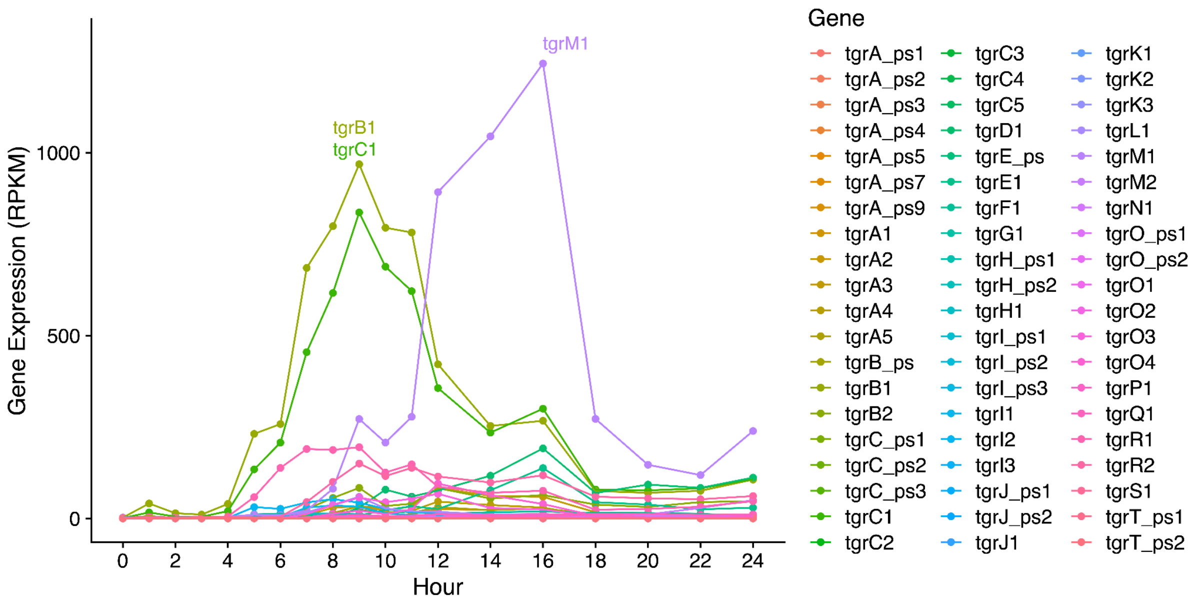

**Figure S3.** PCR assay shows that strains that fail to produce a PCR product for *tgrB2* have a >2kb deletion that impacts the region. (A) Black arrows indicate forward and reverse primers that bind to neighboring genes. (B) Initial assay using strains that do produce tgrB2 products and one strain that does not, revealing a deletion of ~2 kb relative to the other strains. (C) PCR assay applied to a wide range of strains results in 5 strains that lack *tgrB2*: QS18, QS33, QS69, QS73, and QS74, mostly from Texas.

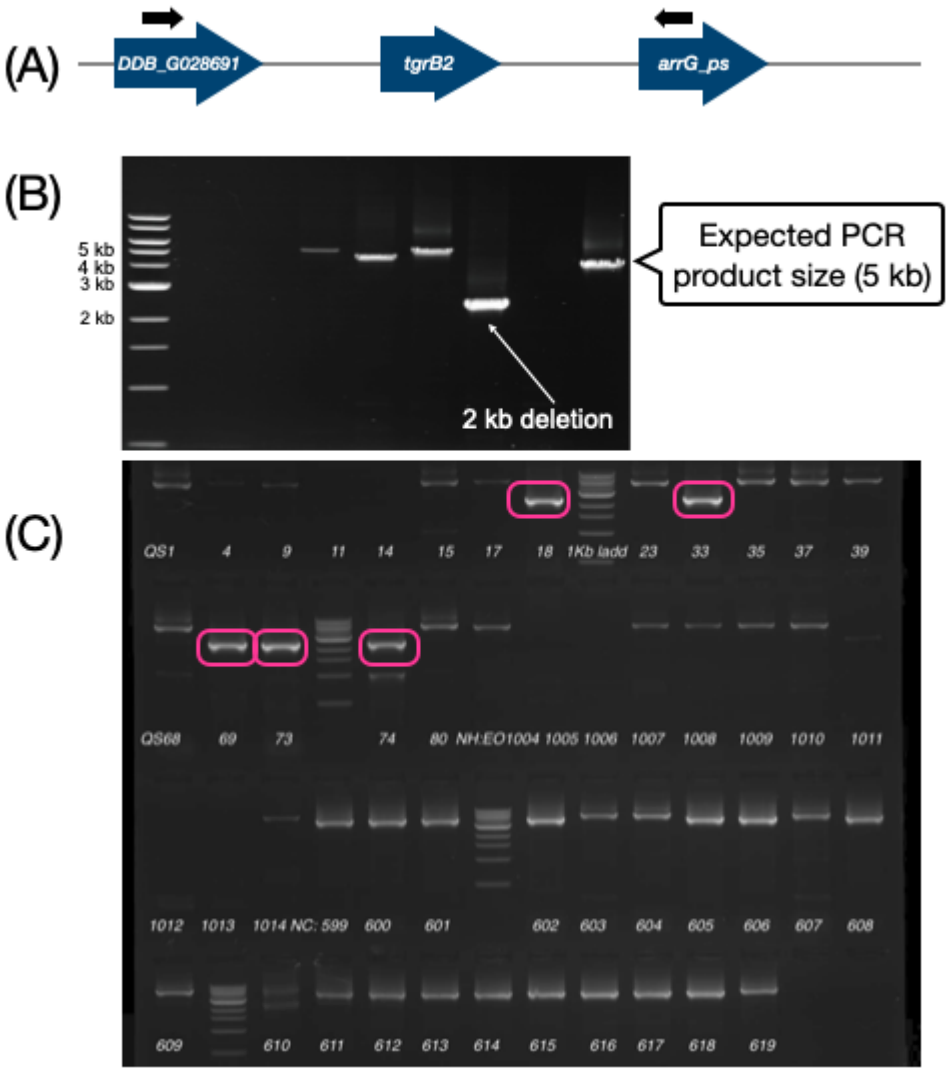

**Figure S4. Pairwise alignment of contigs that harbor the *tgrBC* locus. Alignments were generated using minimap and allowing up to 20% sequence divergence.** (A-C) *D. discoideum* to *D. discoideum* pairs, (D) *D. discoideum* – cryptic pair. Each plot shows alignment of a single contig per strain, and dotted lines indicate the locations of the *tgrBC* genes on each contig. Colored dots indicate the midpoints of an aligned region, which allows very short matches to be visualised. EO1176 has a large insertion compared to QS18, with a breakpoint close to the *tgrBC* locus. Large numbers of dots indicate numerous very short alignments (i.e., highly divergent sequence).

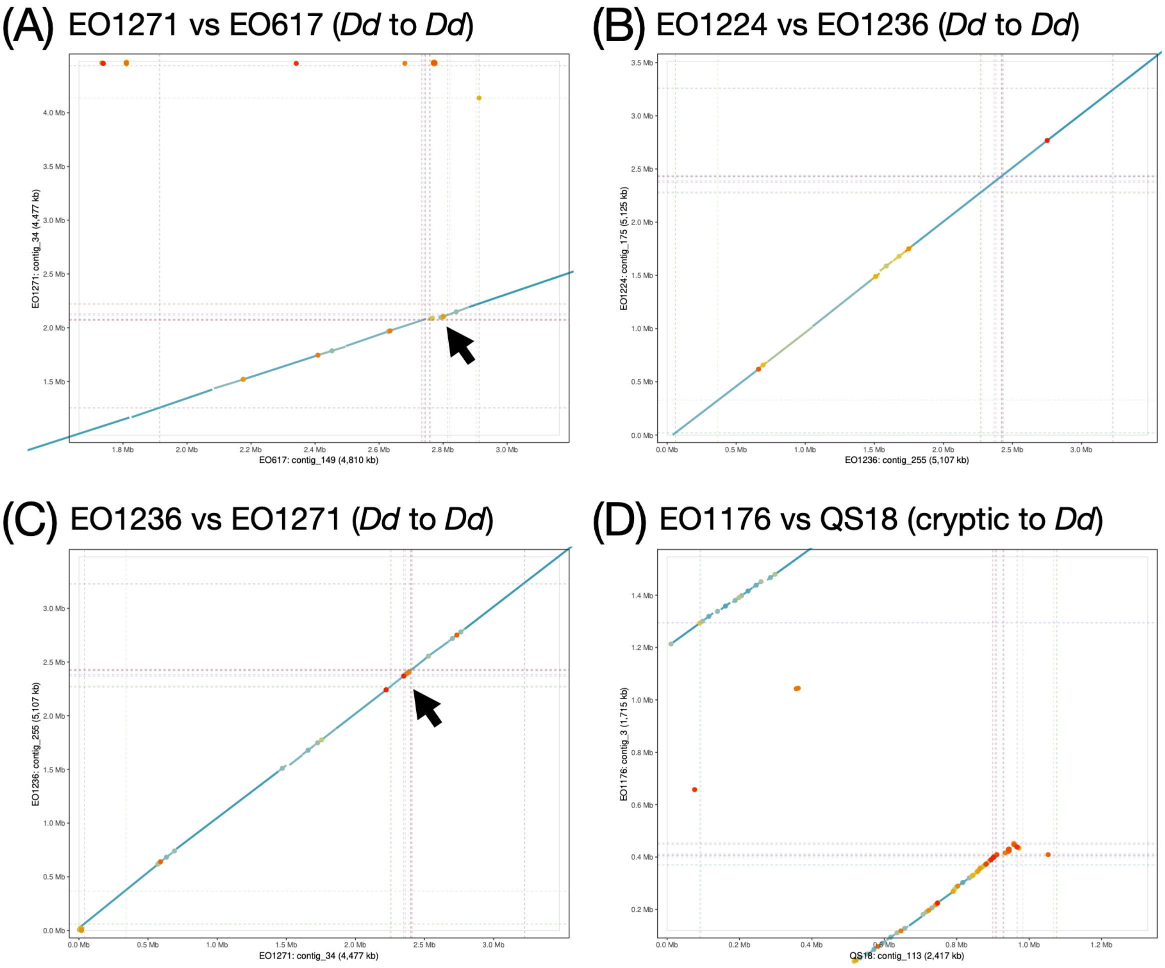

**Figure S5. Pairwise alignment of the *tgrBC* contigs in QS33 (contig_99) and QS68 (contig_48), two *D. discoideum* strains from Texas.** (A) The top contig is QS33 and bottom contig is QS68. Green arrows indicate alignable regions and their corresponding regions in the other contig (connected by light green shading). The two contigs align well over their lengths. The arrow points to the approximate location of the *tgrBC* locus. (B) Same as (A), but zoomed in to the *tgrBC* region. The white area indicates a lack of alignment, corresponding to a ~10kb region in the middle of *tgrBC* locus in QS68 that is absent from QS33, and a smaller region in QS33 that is absent from QS68. (C) Locations of signifcant tblastn hits, using AX4 tgr proteins as queries. Blocks indicate the location of the hit, with the width indicating the size (length) of the hit, and transparency indicating its percent identity. Below each tblastn panel are the locations of the Liftoff-determined *tgr* genes. Liftoff and tblastn agree on the full-length genes, but tblastn finds additional *tgr* gene fragments in the 2.4 kb region.

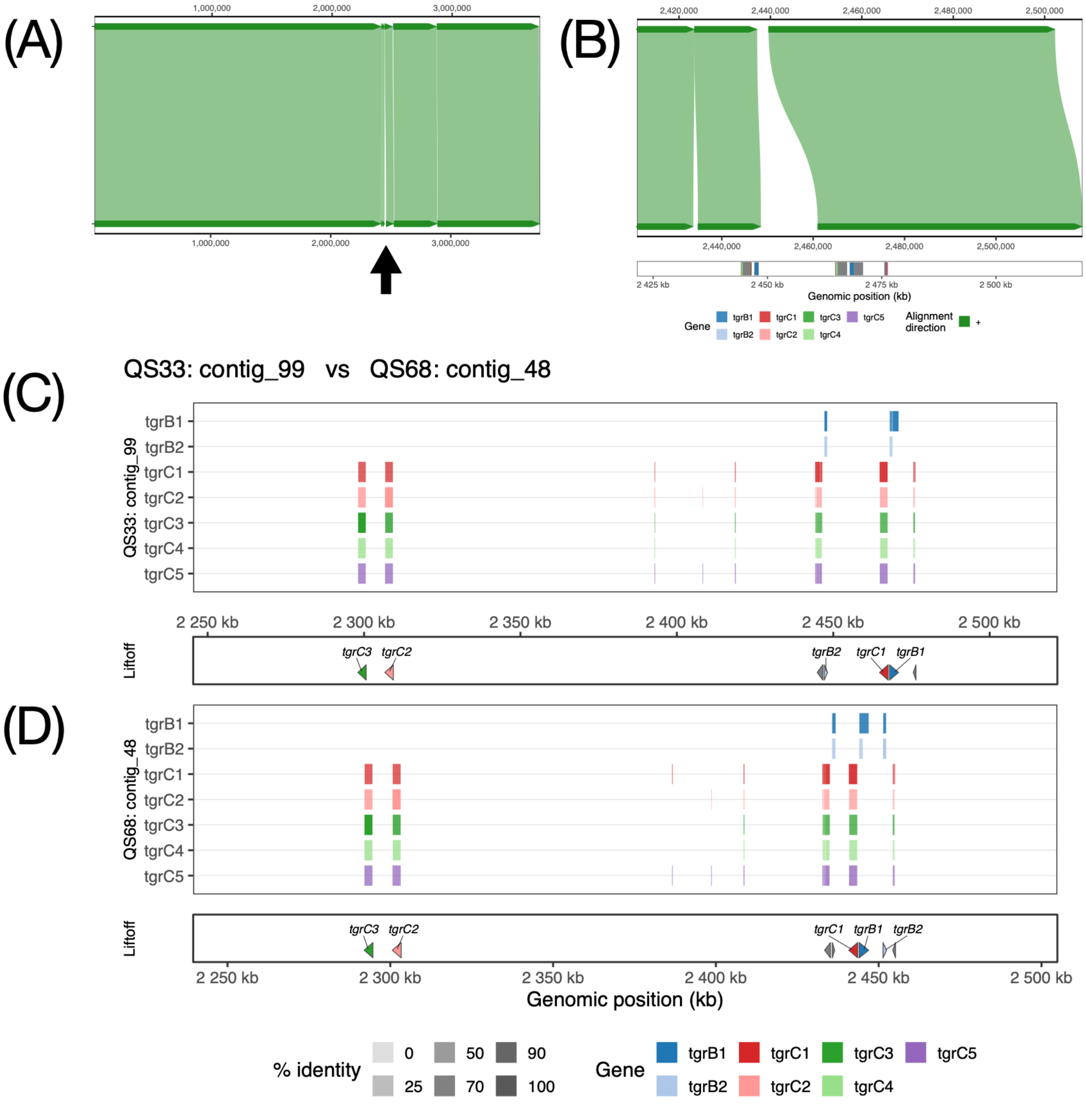

**Figure S6. Spatial analyses of molecular variation in *tgrB1*, *tgrB2*, *tgrC1*, *tgrC2*, *tgrC3*, and *tgrC5*.** Sliding windows for each metric were calculated for each of the four populations in PopGenome and then averaged across populations. Colors indicate locations of protein domains (see Fig. S1).

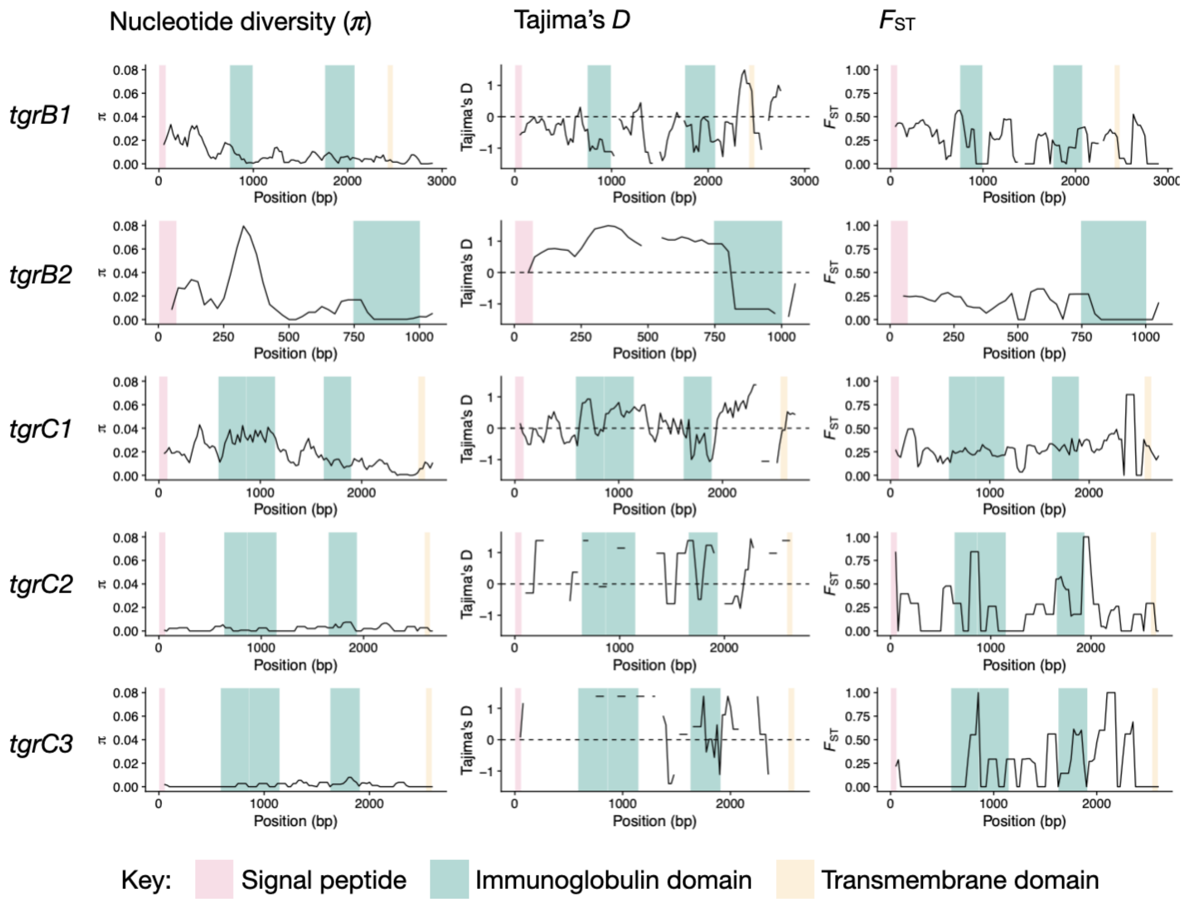

**Figure S7. Mosaic polymorphism in *tgrB2*.** (A) Peak of high nucleotide diversity occurs around base 300, nearly twice the magnitude of the nucleotide diversity observed in *tgrB1* and *tgrC1* (see Fig. S7). (B) Sequence alignment reveals that the boundaries of the polymorphism are highly discrete. (C) Maximum likelihood phylogeny of the three known *tgrB* genes: *tgrB1*, *tgrB2*, and *tgrB* pseudogene in strains sequenced with Nanopore. Phylogeny was rooted using the cryptic species EO1176 as the outgroup. Although the bottom two clades largely correspond to *tgrB1* and *tgrB2* alleles, the top clade consists of a mix of representatives of all three loci, with short branch lengths, indicating recent shared ancestry.

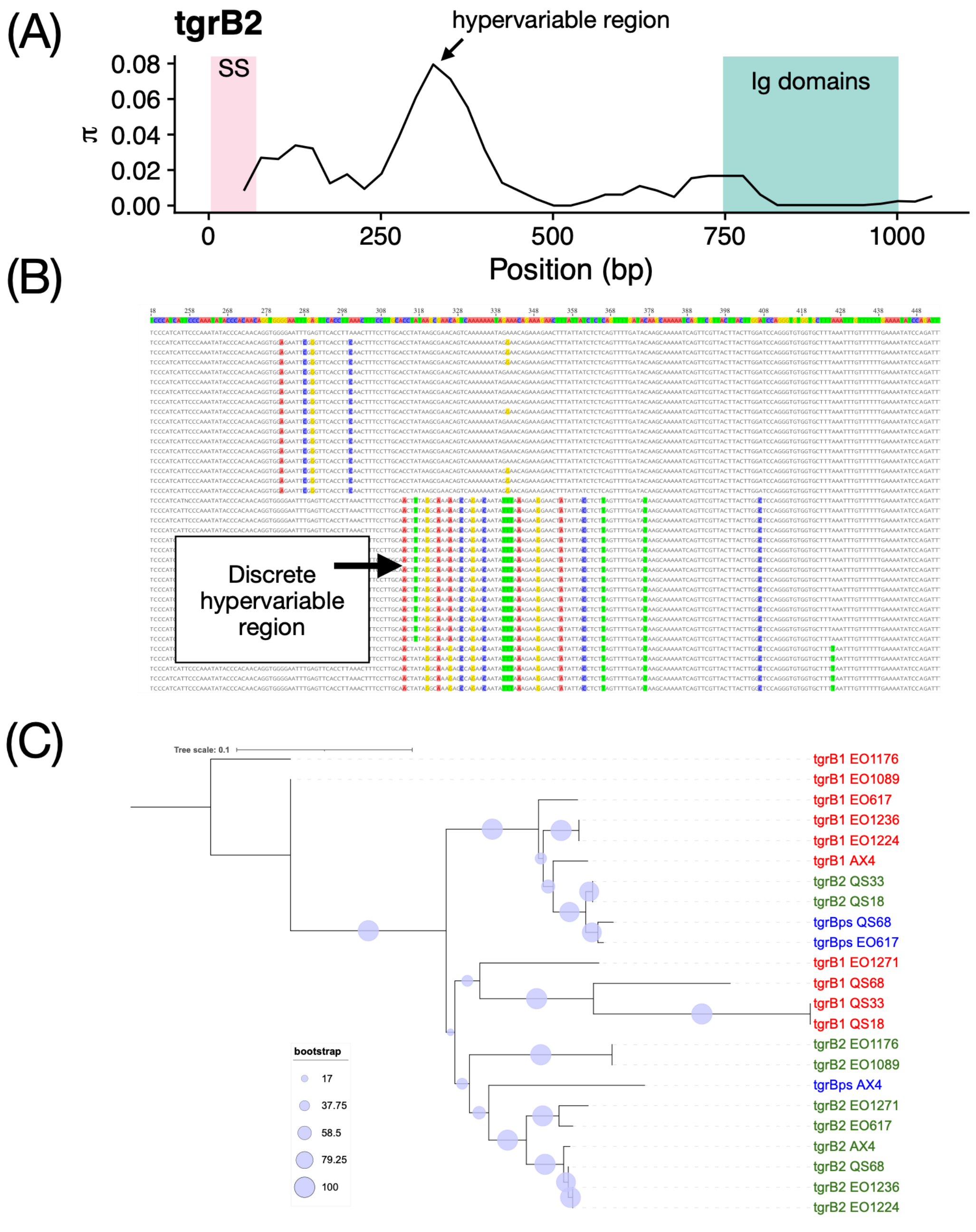

**Figure S8. Intra- and interlocus recombination in the *tgrBC* locus.** (A) Significant recombination detected using RDP. Blue indicates intralocus recombination (but within *tgrB* or *tgrC*), whereas yellow indicates interlocus events (among paralogs). Overlapping hits are overplotted, so darker shading indicates more events in that region. (B) Significant sequence mosaicism using 3Seq. *tgrB* or *tgrC* genes without significant hits are not shown (e.g., *tgrC3*, *tgrC5*). (C) Best estimate of locations for recombination breakpoints using GARD.

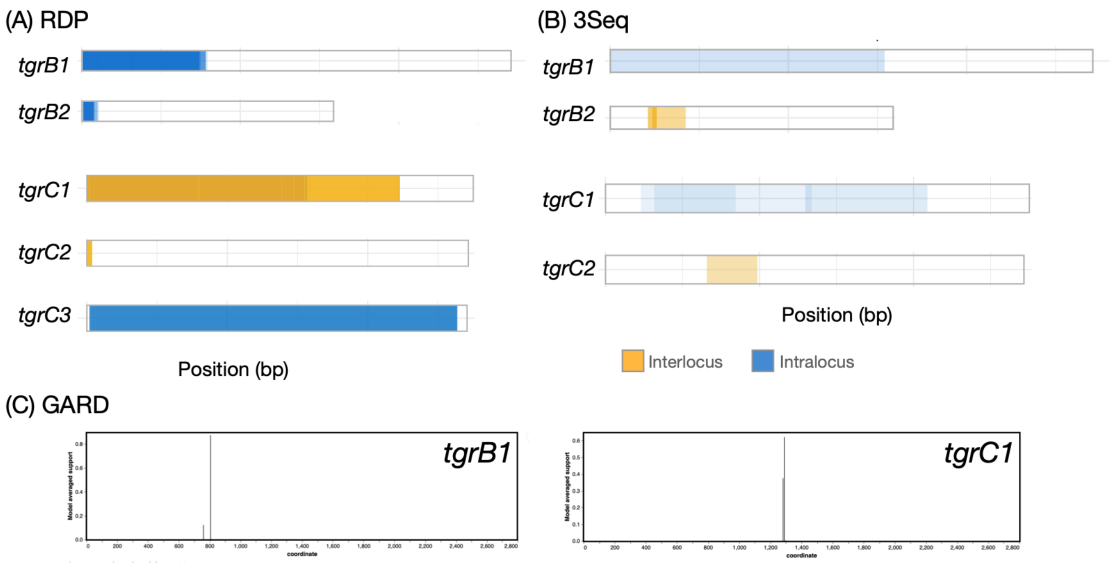

**Figure S9. tblastn results for the *tgrBC* cluster**. The assembly sequences for each strain are anchored on *tgrB1*, which is present in all strains. Vertical dotted lines indicate the start location of the indicated genes in the AX4 reference genome. Colored arrows indicate boundaries of significant tblastn hits using AX4 proteins as the query. Shading indicates the similarity score to AX4, which accounts for the length of the hit, as well as its percent identity (see Methods). This plot matches the Liftoff findings for full-length genes, except for an additional *tgrC* gene in EO1089. Blast also identifies additional gene fragments absent from the Liftoff annotation, which required 100% exon identity to assign as a copy.

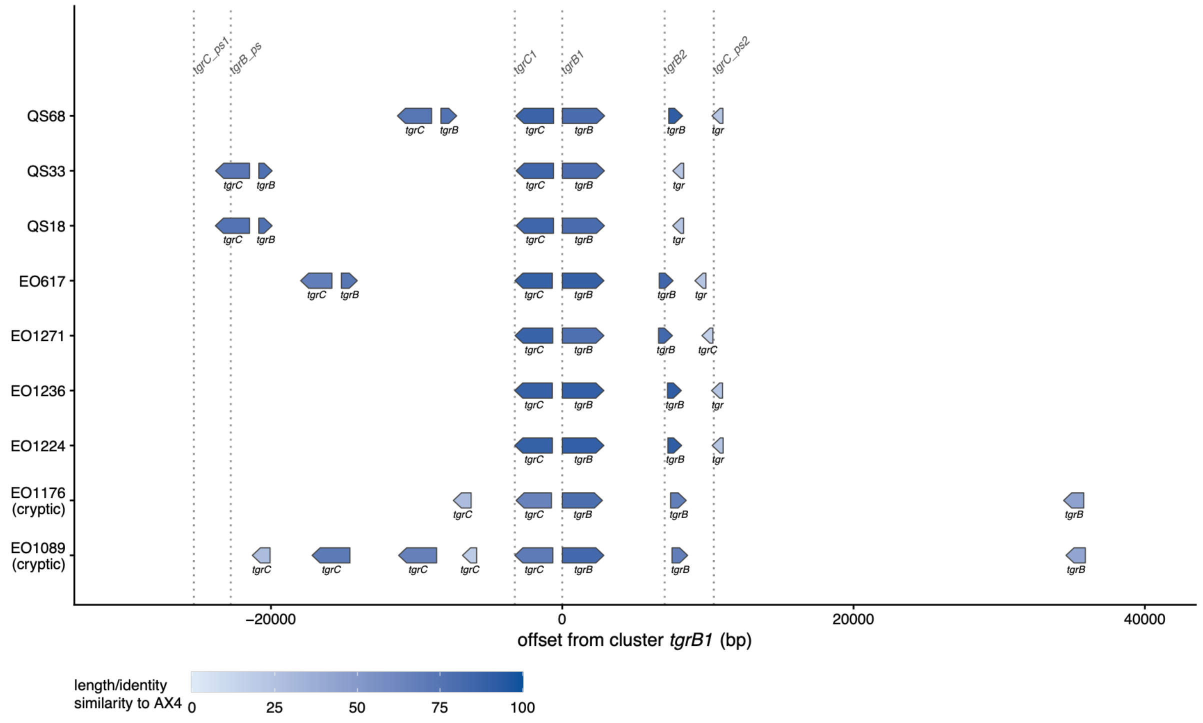

**Figure S10. Length of repeat tract in reads covering EO1176's *tgrC* gene.** Red bars indicate the length of the repeat in the EO1089 (i.e., a non-frameshifted *tgrC* allele), and blue bars indicate the length of the repeat in the EO1176 assembly allele. In four of five cases, EO1176's most common read matches length of the EO1089 allele. In the fifth case (CDS position 2527), the second most common tract length matches the EO1089 allele, and the most common tract length is 12 bases shorter (46 vs 34)—which would indicate a deletion, but not one that results in a frameshift. Overall, there is not strong support for frameshift mutations in EO1176's sole *tgrC* gene.

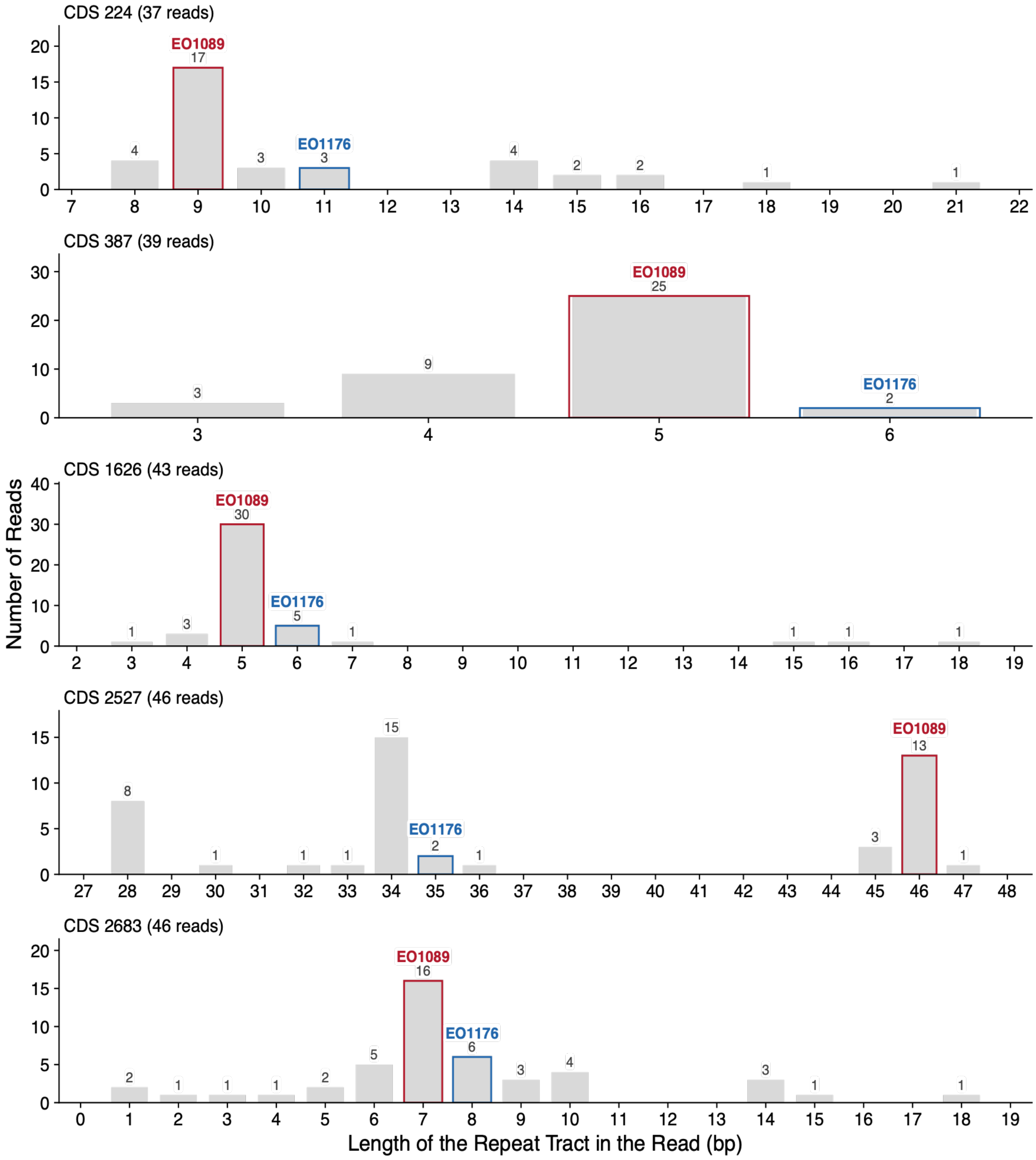
